# Suicidal Thoughts, Behaviors, and Event-Related Potentials: A Systematic Review and Meta-Analysis

**DOI:** 10.1101/2020.04.29.069005

**Authors:** Austin J. Gallyer, Sean P. Dougherty, Kreshnik Burani, Brian J. Albanese, Thomas E. Joiner, Greg Hajcak

## Abstract

**Background:** Suicidal thoughts and behaviors (STBs) are thought to result from, at least in part, abnormalities in various neural systems. Event-related potentials (ERPs) are a useful method for studying neural activity and can be leveraged to study neural deficits related to STBs; however, it is unknown how effective ERPs are at differentiating various STB groups. The present meta-analysis examined how well ERPs can differentiate (a) those with and without suicidal ideation, (b) those with and without suicide attempts, (c) those with different levels of suicide risk, and (d) differences between those with suicide attempts versus those with suicidal ideation only.

**Method:** This meta-analysis included 208 effect sizes from 2,517 participants from 27 studies. We used a random-effects meta-analysis using a restricted maximum likelihood estimator with robust variance estimation. We meta-analyzed ERP-STB combinations that had at least three effect sizes across two or more studies.

**Results:** A qualitative review found that for each ERP and STB combination, the literature is highly mixed. Our meta-analyses largely did not find significant relationships between STBs and ERPs. We also found that the literature is likely severely underpowered, with most studies only being sufficiently powered to detect unrealistically large effect sizes.

**Conclusions:** Our results provided little-to-no support for a reliable relationship between the ERPs assessed and STBs. However, the current literature is severely underpowered, and there are many methodological weaknesses that must be resolved before making this determination. We recommend large-scale collaboration and improvements in measurement practices to combat the issues in this literature.

---

Suicide is a concern around the world. Estimates indicate that well over 800,000 people die by suicide annually (World Health Organization, 2014). In the United States alone, over 47,000 people die by suicide each year, making suicide the 10^th^ leading cause of death (Centers for Disease Control, 2017). In contrast to trends in other western countries, suicide rates in the United States have been increasing over the last two decades (Hedegaard et al., 2018). This increase in the suicide rate has contributed to the decreasing trend in U.S. life expectancy every year since 2014 (Woolf & Schoomaker, 2019).

Many scientists have suggested that suicidal thoughts (i.e., thoughts related to desire for death or suicide, regardless of suicidal intent) and behaviors (i.e., suicide attempt with nonzero intent; death by suicide; Silverman et al., 2007) are partially the result of differences or abnormalities in neurobiological systems (Joiner et al., 2005; Mann, 2003; Van Heeringen & Mann, 2014). Specifically, most of these neurobiological models of suicide suggest that suicide is the result of an interaction between dynamic, contextual factors (e.g., stress) and static factors (e.g., genetic loading for suicidal behavior that is independent of mental disorders; Mann, 2003, 2003; Mann & Rizk, 2020; Van Heeringen & Mann, 2014). Variations in genetic loading, these models argue, play an important role in the development of the structure and function of neural circuits, such as in the hypothalamic-pituitary-adrenal axis or in the serotonergic projections throughout the brain (Mann & Rizk, 2020). This diathesis-stress model has led to studies investigating the brain regions that are predicted to be implicated in suicidal thoughts and behaviors (STBs), with most studies using magnetic resonance imaging (MRI) to examine structural and functional brain differences between those with and without a history of STBs. These findings, however, have been mixed, as illustrated by two recent, competing reviews of this literature.

The first, a qualitative, systematic review of the neuroimaging literature of STBs, concluded that there is tentative but converging evidence for two brain networks implicated in STBs (Schmaal et al., 2019). First, the authors proposed that the ventral prefrontal cortex and many of its connections are involved in increasing negative and decreasing positive internal states, which may lead to suicidal ideation (SI). Second, the authors identified other regions, including the dorsomedial prefrontal cortex, the dorsolateral prefrontal cortex, and the inferior frontal gyrus, as a separate network that may be important for suicide attempts (SAs) due to these regions’ roles in planning and cognitive control. In contrast, a quantitative meta-analysis of neuroimaging studies using whole-brain analyses did not find significant associations between SAs and any brain region; the study concluded that there were not enough studies to analyze the relationship between SI and neural structure and function (Huang et al., 2020). The mixed evidence illustrated by these two reviews underscores the need to broaden the methods used to examine differences in neural functioning among those experiencing STBs.

An alternative method to examining the neurobiology of STBs that has been gaining more attention is to use event-related potentials (ERPs). ERPs are derived from electroencephalographic (EEG) data and reflect neural responses to specific, repeated events (Luck, 2014). ERP waveforms are calculated by averaging the electrocortical responses to a specific task event (e.g., stimulus presentation, error commission) across multiple trials. Although ERPs generally lack the excellent spatial resolution of MRI (i.e., the ability to accurately localize neural functions to regions in the brain), this methodology has several strengths. First, ERPs are direct measures of brain activity with excellent temporal resolution, which allows researchers to examine distinct neural activity that might typically overlap when assessed in MRI (e.g., ERPs can separately index multiple distinct processes that occur within hundreds of milliseconds of one another). Many ERPs have been shown to have excellent internal consistency with sufficient trial numbers (e.g., Meyer et al., 2013; Moran et al., 2013). Electrocortical data are also less expensive to collect with fewer counterindications (e.g., metal in body; claustrophobia), relative to other forms of neuroimaging, thereby making larger samples sizes more feasible. Importantly, the extant literature has demonstrated that many ERPs reflect different neural functions—such as emotion regulation, response inhibition, reward processing, reward anticipation, and error monitoring (Luck & Kappenman, 2011)—that might be relevant for understanding STBs.

Because of their potential clinical utility (Hajcak et al., 2019), researchers have begun to employ ERPs in the study of STBs. For example, Gibb and Tsypes (2019) suggested that the inclusion of ERPs in machine learning algorithms may improve the prediction of STBs. Though there have been studies examining differences in ERPs between those with SI or a history of SAs and those without SI or SAs (e.g., Albanese et al., 2019a), the literature on the relationship between ERPs and STBs is mixed. As an example, we will briefly discuss the late positive potential (LPP)in relation to STBs.

The LPP is a sustained positive deflection during the presentation of emotionally valanced stimuli, with the LPP demonstrating greater amplitudes in response to emotional stimuli (e.g., images of mutilation and sex), relative to neutral stimuli (Cuthbert et al., 2000; Weinberg & Hajcak, 2010, 2011). The LPP has been shown to correlate with activity in cortical and subcortical areas—including areas such as the amygdala, ventrolateral prefrontal cortex, posterior cingulate cortex, and orbitofrontal cortex—that form an affective-salience network (Liu et al., 2012). Importantly, the blunting of the LPP in response to threatening stimuli has been hypothesized to index fearlessness about death, a construct that leading theories of suicide posit is key for individuals to overcome the instinct of self-preservation and enact lethal self-injury (Gallyer, Hajcak, et al., 2020; Klonsky & May, 2015; O’Connor & Kirtley, 2018; Van Orden et al., 2010). That is, no matter how much a person may want to die by suicide, they will be unable to do so unless they also have a reduced fear response to the prospect of their own death. Though more research is still needed, preliminary evidence has shown that the LPP may be related to self-reported fearlessness about death (Bauer et al., 2020). Thus, the LPP is a neural measure of affective processing that may be important for the transition from (a) thinking about suicide to (b) engaging in suicidal behaviors. Consistent with this possibility, one study found that the LPP to threatening stimuli was blunted in patients with a previous SA, compared to patients without a previous SA (Weinberg et al., 2017). However, the evidence of the relationship between the LPP and SI has been more mixed. One study found that a reduced LPP to pleasant pictures was related to increased SI (Weinberg et al., 2016). In contrast, a follow-up study did not find a relationship between SI and the LPP to rewarding stimuli (Weinberg et al., 2017), and another study found such a relationship only when individuals were asked to volitionally increase their reward responses (Albanese et al., 2019b).

Though there are many reasons for these mixed findings, one possible explanation involves the definitions of STBs used across studies. According to ideation-to-action theories of suicide (e.g., interpersonal theory of suicide; 3-step theory; Joiner, 2005; Klonsky & May, 2015), factors that predict SI do not necessarily predict SAs or death by suicide (Klonsky et al., 2018). This finding has been supported by a meta-analysis showing that most risk factors for suicide (e.g., depression) have stronger associations with SI than with SAs (May & Klonsky, 2016). Given these findings, some ERPs may relate to SI, whereas others may relate to SAs or to other STB outcomes, depending on the neural process indexed by specific ERPs. For example, we previously mentioned that the LPP to threatening stimuli may be a measure of fearlessness about death, which may be most relevant for the transition from suicidal thoughts to engaging in suicidal behavior. In contrast, the blunting of a separate ERP, called the reward positivity (RewP), may be more likely to be related to SI, rather than to SAs, given the RewP’s well-established relationship to depressive symptoms (Keren et al., 2018).

In addition to the above mixed findings, recent meta-analyses have called into question the clinical significance of most risk factors for STBs when examined in isolation (Chang et al., 2016; Franklin et al., 2017; Ribeiro et al., 2018). Researchers have also called into question whether the modal study across neural methods (e.g., fMRI and ERPs) is sufficiently powered to detect individual differences (Clayson et al., 2019; Elliott et al., 2020; Marek et al., 2020). Thus, a meta-analysis of all ERPs is needed to summarize the current literature’s mixed findings, and to provide a best estimate of the true effect size of the relationship between STBs and ERPs so that researchers in this burgeoning area can ensure that they are using designs that are adequately powered. Moreover, given that this research area is gaining momentum, it is important to qualitatively review the current literature for individual ERPs and their relationships to STBs. Thus, the current systematic review and meta-analysis aims to: (1) review the literature to examine how various ERPs are related to STBs; (2) provide meta-analytic estimates of whether any ERPs differ between individuals who have and have not engaged in STBs; and (3) examine how powered the literature is across a range of effect sizes. We examined each of these aims in relation to four separate STBs: (1) between those with and without suicidal ideation, (2) between those with and without a suicide attempt history, (3) among those with varying degrees of suicide risk, and (4) between those with a previous suicide attempt and those with suicidal ideation but without a previous suicide attempt. Notably, we did not examine any moderators in our meta-analyses for two reasons: (1) many ERP-STB combinations have relatively few studies, making moderation analyses significantly underpowered and (2) each ERP may have several moderators that are unique to that ERP (e.g., if the RewP effect moderated by nature of the reward, such as rewarding pictures or money). Given that our data is freely available (see Method section), we recommend that future investigators conduct meta-analyses focused on one ERP at a time and explore whether there may be any moderators for these meta-analytic effects. Based on the existing risk factor literature of STBs, we hypothesized that ERPs would have a small-to-moderate relationship with STBs. We did not have any specific hypotheses for our other aims.

## Method

### Inclusion and Exclusion Criteria

All of our data, analysis scripts, coding spreadsheets, figures, and supplementary material are available on the Open Science Framework (https://osf.io/k4cpe/?view_only=612a0952f8834206813d8414a7018606). Our meta-analysis was conducted in accordance with the PRISMA Statement (Moher et al., 2009). The following criteria were established to select relevant effect sizes.

#### Language

Only articles written in English were included.

#### Event-Related Potentials

Only articles that had an effect size that included an event-related potential (ERP) were included. ERPs were defined as measures using EEG responses to specific, time-locked events of mental processes. Given these criteria, studies that used cortical recordings or that stimulated the brain to produce neural activity were excluded.

#### Suicide-Related Outcomes

Effect sizes were required to include a measure of suicidal thoughts, suicidal behaviors, or suicide risk/suicidality. Measures of suicidal thoughts were defined as any measure of severity or history of suicidal ideation. We did not distinguish between death/suicidal ideation or between passive/active suicidal ideation. Suicidal behaviors were defined as a history of engaging in a self-harm behavior with nonzero intent to die. Importantly, we did not distinguish among aborted, interrupted, and actual SAs. Moreover, effect sizes that included SA frequency were not included. Suicide risk was included as an umbrella term when studies did not cleanly delineate between individuals with a history of SAs and those with a history of SI but no history of SAs. For example, studies that used the Suicidal Behavior Questionnaire—Revised (SBQ-R; Osman et al., 2001) were categorized as “suicide risk” studies because this measure asks questions about suicidal thoughts and behaviors, and gives a sum score. Importantly, the measures in studies that were coded as “suicide risk” could be either dimensional (e.g., when the measure was the SBQ-R) or categorical (e.g., group was the “suicidal group” if they had history of SI or a history of SA). Given these definitions, studies that examined nonsuicidal self-injury or a history of nonsuicidal self-injury were excluded. Last, we compared studies that included a group that had a history of SAs, and a group that had a history of SI but no history of SAs. These four outcome differences were chosen to reflect the current ideation-to-action framework that guides most suicide research.

#### Published in Print or Online by October 2020

Our search results were limited to articles published in print or online by October 20, 2020.

### Literature Search

We conducted literature searches across PubMed, PsycInfo, Web of Science, and ProQuest Dissertations & Theses to find relevant literature. Full search terms for each database are available on an online repository stored on the Open Science Framework https://osf.io/k4cpe/?view_only=612a0952f8834206813d8414a7018606). To summarize, search terms included different permutations related to ERPs and suicidal thoughts and behaviors, including: “event-related potential,” “event-related potentials,” “electroencephalography,” “EEG,” “evoked potential,” “evoked potentials,” “late positive potential,” “reward positivity,” “P3a,” “P3b,” “error-related negativity,” “suicide,” “suicidal,” “suicidal ideation,” “self-harm,” “self-injury,” “parasuicide,” “suicide ideation,” “suicide attempt,” “suicide plan,” “suicidality,” “suicide risk,” “attempter,” “attempters,” and “ideators.” We also checked the reference list of every article included in the final meta-analysis for additional studies. In addition to these efforts, we made several attempts to gather unpublished literature. These efforts included: (1) social media announcements on Twitter and (2) contacting researchers who were either a first or last author on at least two papers included in the meta-analysis. Of the nine researchers we emailed asking for unpublished or yet-to-be-published studies, six responded, with three providing data (two of which jointly contributed a dataset) and three indicating that they did not have any other unpublished data. The remaining three did not respond. We also had one unpublished paper that was included in this meta-analysis. We first conducted this literature search in October 2019 but then updated the literature search in October 2020 because enough time had elapsed to make the systematic review and meta-analysis outdated. For the updated literature search, we used the same process, except we limited the dates on all our databases to only show articles published after the date of our initial literature search in October 2019.

### Data Extraction and Coding

For each effect size, the following were also coded: mean age of the sample, suicide measure, EEG reference, scoring ERP procedure, specific ERP, type of suicidal group, and the type of comparison group. All studies were initially coded by the first author (A.J.G.) in an Excel spreadsheet. One co-author (S.P.D.) recoded data included in the meta-analysis. We conducted reliability analyses for our codings, using agreement rate and Cohen’s kappa for categorical variables and using intercoder correlation and two-way random-effects intraclass correlation (ICC) for continuous variables. For categorical variables, agreement rate ranged from 88.0% to 100%, and Cohen’s kappas were excellent (κ = .84–.99). For continuous variables, intercoder correlations ranged from .99 to 1.0, and ICCs also ranged from .99 to 1.0. Disagreements for categorical variables were resolved through discussion.

Using our search terms across all our included databases produced 298 papers. Figure 1 displays an overview of our search process and the number of articles within each search process. After accounting for duplicates, we examined the abstracts of 215 papers and excluded studies that did not meet inclusion criteria. After examining abstracts, 51 studies entered full-text review. After full-text review of the literature from our database search, 23 studies were included. Examining citations from these papers added one more study, and our efforts to reach out to authors for unpublished data resulted in three more studies. In the end, 27 studies had at least one effect size that met inclusion criteria, resulting in 208 total effect sizes. The 29 studies that were not included after full-text review (including one study we found through examining citations) were excluded for the following reasons: (1) no measure that met our definition of ERP (*n* = 3), (2) no measure that met one of our definitions of STBs (*n* = 7), (3) no or insufficient information to calculate effect sizes, including when access to the full text was inaccessible due to the article’s being a poster or dissertation (*n* = 17), (4) paper was a review article (*n* = 1), and (5) effect size from paper was a duplicate of that from another paper (*n* = 1). For articles for which the full text was available but for which we did not have enough information to calculate effect sizes, we sought to email the corresponding or last author. We emailed four researchers for more information to calculate effect sizes from their papers. One responded and indicated that they no longer had access to the data. No other researchers responded, and we were unable to locate contact information for authors for two studies. See Table 1 for included studies.

**Figure 1.**
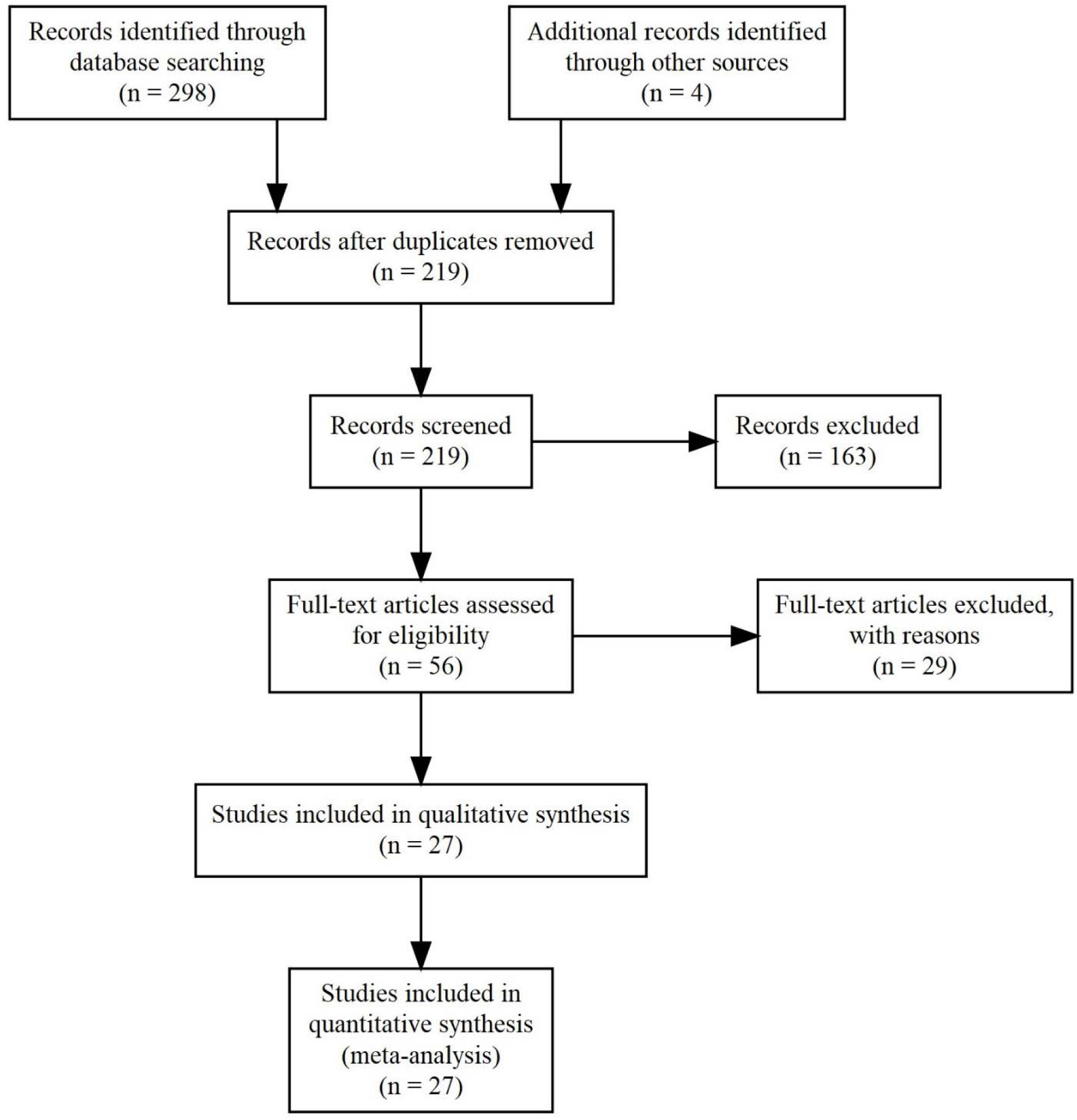
PRISMA Diagram

**Table 1.**
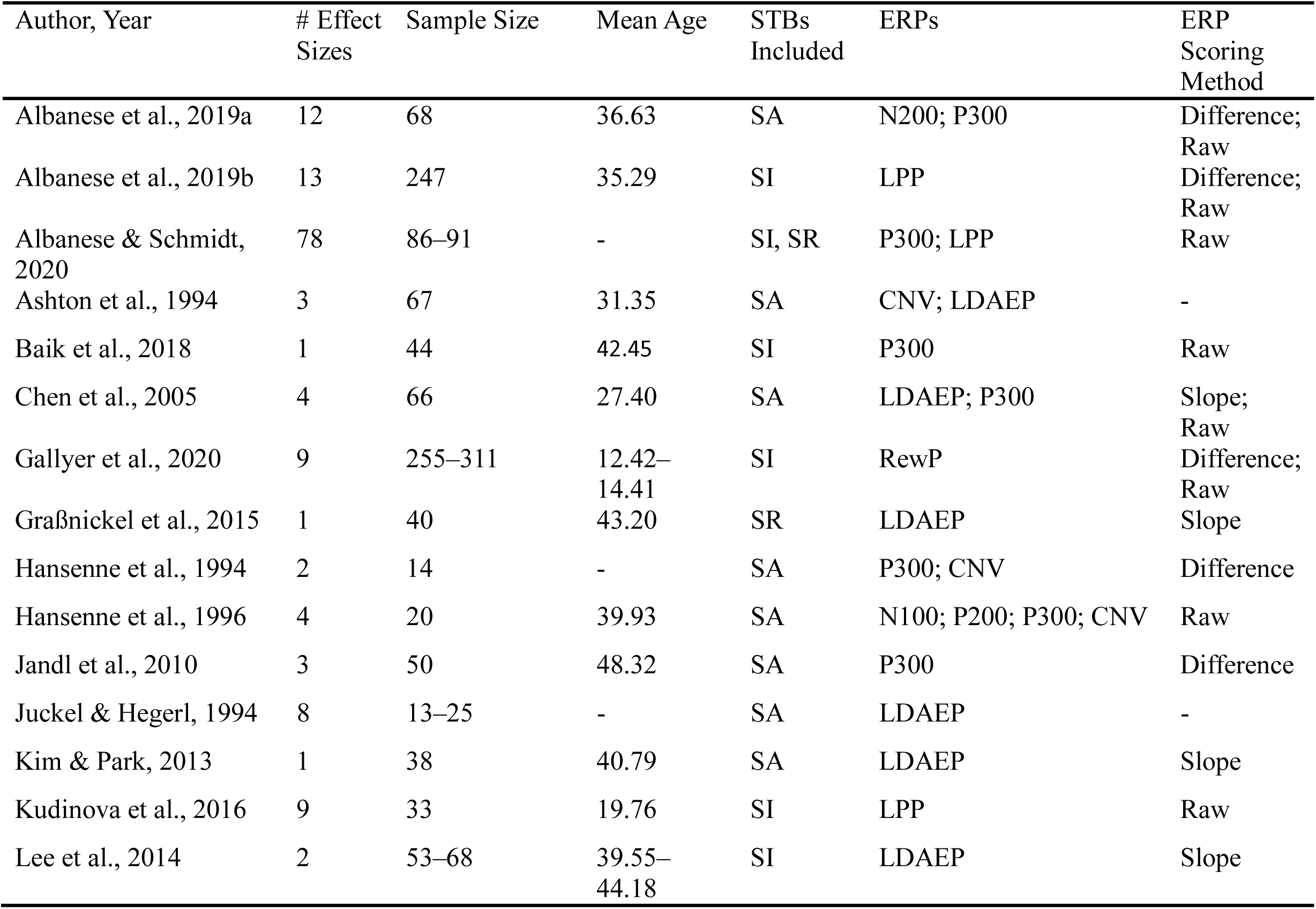
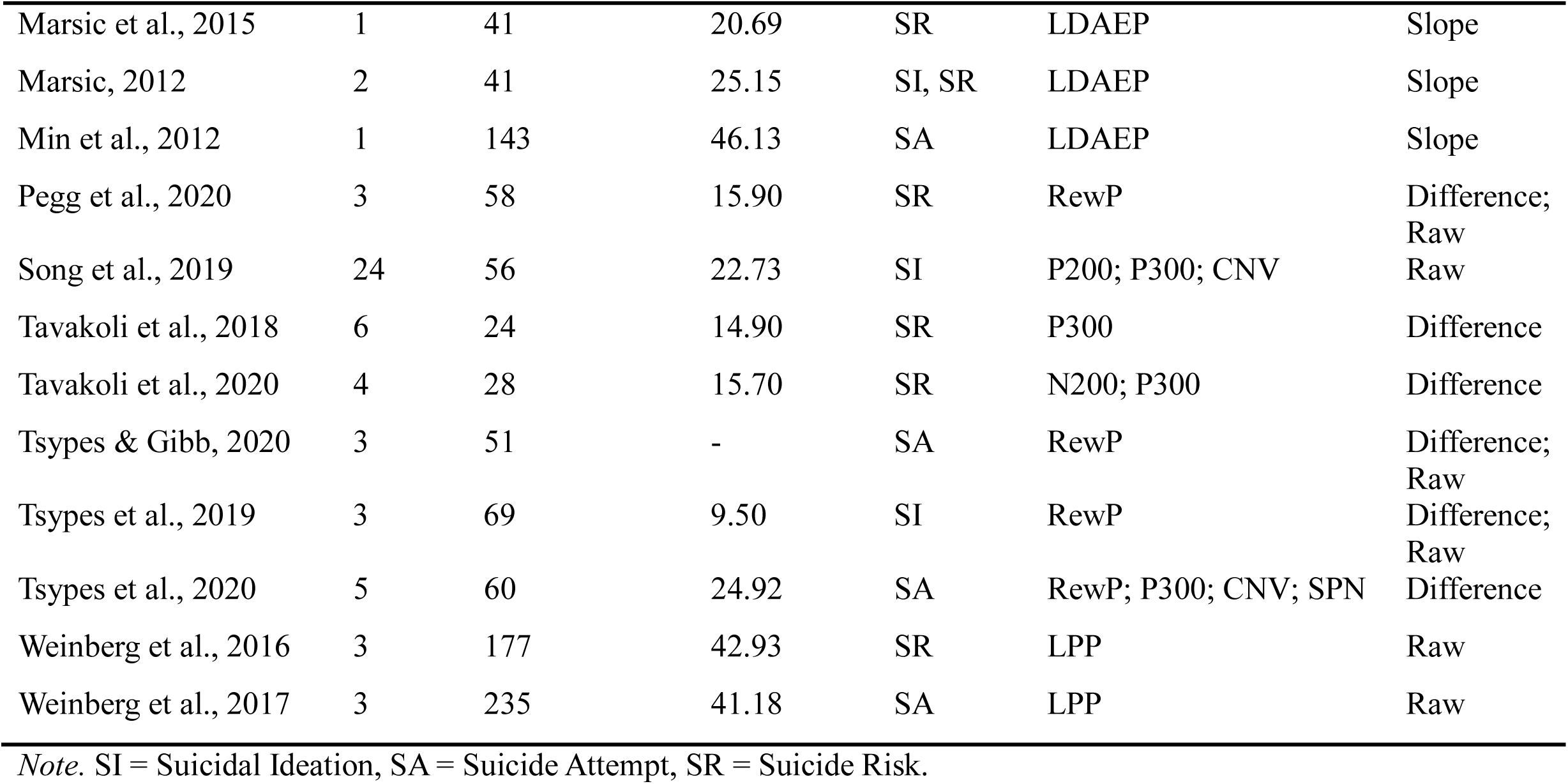
Included Studies and Study Characteristics

| Author, Year | # Effect Sizes | Sample Size | Mean Age | STBs Included | ERPs | ERP Scoring Method |
| --- | --- | --- | --- | --- | --- | --- |
| Albanese et al., 2019a | 12 | 68 | 36.63 | SA | N200; P300 | Difference; Raw |
| Albanese et al., 2019b | 13 | 247 | 35.29 | SI | LPP | Difference; Raw |
| Albanese & Schmidt, 2020 | 78 | 86–91 | - | SI, SR | P300; LPP | Raw |
| Ashton et al., 1994 | 3 | 67 | 31.35 | SA | CNV; LDAEP | - |
| Baik et al., 2018 | 1 | 44 | 42.45 | SI | P300 | Raw |
| Chen et al., 2005 | 4 | 66 | 27.40 | SA | LDAEP; P300 | Slope; Raw |
| Gallyer et al., 2020 | 9 | 255–311 | 12.42–14.41 | SI | RewP | Difference; Raw |
| Graßnickel et al., 2015 | 1 | 40 | 43.20 | SR | LDAEP | Slope |
| Hansenne et al., 1994 | 2 | 14 | - | SA | P300; CNV | Difference |
| Hansenne et al., 1996 | 4 | 20 | 39.93 | SA | N100; P200; P300; CNV | Raw |
| Jandl et al., 2010 | 3 | 50 | 48.32 | SA | P300 | Difference |
| Juckel & Hegerl, 1994 | 8 | 13–25 | - | SA | LDAEP | - |
| Kim & Park, 2013 | 1 | 38 | 40.79 | SA | LDAEP | Slope |
| Kudinova et al., 2016 | 9 | 33 | 19.76 | SI | LPP | Raw |
| Lee et al., 2014 | 2 | 53–68 | 39.55–44.18 | SI | LDAEP | Slope |
| Marsic et al., 2015 | 1 | 41 | 20.69 | SR | LDAEP | Slope |
| Marsic, 2012 | 2 | 41 | 25.15 | SI, SR | LDAEP | Slope |
| Min et al., 2012 | 1 | 143 | 46.13 | SA | LDAEP | Slope |
| Pegg et al., 2020 | 3 | 58 | 15.90 | SR | RewP | Difference;<br>Raw |
| Song et al., 2019 | 24 | 56 | 22.73 | SI | P200; P300; CNV | Raw |
| Tavakoli et al., 2018 | 6 | 24 | 14.90 | SR | P300 | Difference |
| Tavakoli et al., 2020 | 4 | 28 | 15.70 | SR | N200; P300 | Difference |
| Tsypes & Gibb, 2020 | 3 | 51 | - | SA | RewP | Difference;<br>Raw |
| Tsypes et al., 2019 | 3 | 69 | 9.50 | SI | RewP | Difference;<br>Raw |
| Tsypes et al., 2020 | 5 | 60 | 24.92 | SA | RewP; P300; CNV; SPN | Difference |
| Weinberg et al., 2016 | 3 | 177 | 42.93 | SR | LPP | Raw |
| Weinberg et al., 2017 | 3 | 235 | 41.18 | SA | LPP | Raw |

*Note.* SI = Suicidal Ideation, SA = Suicide Attempt, SR = Suicide Risk.

### Effect Sizes

Information was extracted from each study to calculate Hedges’ *g*. For most effect sizes, means, standard deviations, and sample size were used to calculate effect sizes. However, in some cases, existing effect sizes had to be converted to Hedges’ *g*. Specifically, several studies reported Pearson’s *r* or an *F* value. For one effect size, Pearson’s *r* was estimated by using the computer program WebPlotDigitizer (Version 4.2; Rohatgi, 2019). All effect sizes were calculated using the “compute.es” package (Version 0.2-4; Del Re, 2013). For all our analyses, a positive Hedges’ *g* means the ERP is larger in those with STBs (e.g., SI) compared to controls, whereas a negative Hedges’ *g* means the ERP is smaller in those with STBs.

### Combining Effect Sizes

We used R (Version 4.0.2; R Core Team, 2020) to conduct all of our analyses and create all of our figures using the following packages: “metafor” (Version 2.4; Viechtbauer, 2010), “tidyverse” (Version 1.3.0; Wickham et al., 2019), “readxl” (Version 1.3.1; Wickham & Bryan, 2019), “PRISMAstatement” (Version 1.1.1; Wasey, 2019), “metaviz” (Version 0.3.1; Kossmeier et al., 2020), “metameta” (Version 0.1.1; Quintana, 2020a), and “ggthemes” (Version 4.2.0; Arnold, 2019). We opted to use a random-effects model that estimated heterogeneity using restricted maximum likelihood. We chose this model because our literature search produced studies with many effect sizes. This dependence between effect sizes violates the assumptions of fixed-effects models, which would lead to an incorrectly estimated pooled effect size and would increase the probability of a Type I error (Hedges et al., 2010). Our random-effects model included two levels: (1) effect size and (2) journal article. We also used robust variance estimation to account for the small number of clusters (i.e., studies) in our analyses (Hedges et al., 2010). ERPs that had at least three effect sizes within a STB outcome (e.g., RewP and SI) and consisted of at least two separate studies, were meta-analyzed.

### Estimating Publication Bias, and Statistical Power

For ERP-STB combinations with at least ten effect sizes, we examined whether publication bias or small-study bias was present by conducting a variant of Egger’s regression for detecting funnel plot asymmetry, using the square root of the effect size weight as the predictor (i.e., √W_i_). Simulation studies have found that this approach balances the Type I error rate (i.e., detecting funnel plot asymmetry when there is none) and statistical power in the presence of dependent effect sizes and when using standardized means difference effect sizes (Pustejovsky & Rodgers, 2019; Rodgers & Pustejovsky, 2020). Finally, to estimate the statistical power of each study for a range of true effect sizes, we used the “metameta” package and adapted code from a freely available repository (Quintana, 2020b). Specifically, for each effect size within each study we calculated the statistical power for ten true effect sizes, ranging from 0.1 to 1, increasing in increments of 0.1. Next, within each study we used the effect size with the median power for all analyses and visualizations.

## Results

Our literature search resulted in 27 studies meeting inclusion criteria, with the earliest included studies being published in 1994 and with a rising number of papers being published every year since 2017. Our results will be presented as follows: (1) a narrative review of each ERP and its relation to individual STBs; (2) for ERP-STB combinations with at least three effect sizes across two studies, a summary of meta-analysis and publication bias results; and (3) statistical power across all studies given a range of effect sizes.

### Narrative Review

#### Loudness-Dependent Auditory Evoked Potential (LDAEP)

The loudness-dependent auditory evoked potential (LDAEP) is an ERP that is calculated as the amplitude change of the N1/P2 component that arises in response to increasingly intense auditory stimuli (Hegerl et al., 2001). The LDAEP has been shown to be negatively related to serotonergic activity in the central nervous system and has been proposed as a reliable indicator of serotonin levels in humans (Hegerl & Juckel, 1993; Juckel et al., 1997). To date, only two studies have examined the LDAEP and how it may differ in those with SI. The first, a study of college students and community members, did not find a relationship between the LDAEP and self-reported SI (*g* = 0.31; Marsic, 2012). The second study recruited two groups of patients: (1) patients with depression with atypical features and (2) patients with depression without atypical features (Lee et al., 2014). The study found a significant negative relationship between the LDAEP and self-reported SI but only in patients with atypical depression (*g* = - 0.83); there was no evidence of a relationship between the LDAEP and SI in patients with typical depression (*g* = -0.01). Though this pattern suggests a potential interaction between LDAEP and atypical versus typical depression, this study did not formally test this interaction, so it is an open question whether the LDAEP—and, in turn, central serotonergic activity—is related to SI in patients with atypical depression.

The literature examining differences in the LDAEP between those with SA versus those without a previous SA is highly mixed. In three independent samples, Juckel and Hegerl (1994) found that those with a previous SA had a smaller LDAEP, relative to controls (*g* = -1.52 – -0.33). Though another study found similar results, other studies have yielded opposing or null findings (*g* = -0.52; Ashton et al., 1994). For example, a study in patients with depression did not find differences in the LDAEP between patients with a previous SA and patients with no previous SA (*g* = 0.65; Chen et al., 2005). Another study similarly did not find any differences between those with a SA and those without (*g* = 0.16; Min et al., 2012). Yet another study among patients diagnosed with depression found that patients with a previous SA had a larger LDAEP, compared to patients without a previous SA (*g* = 0.69; Kim & Park, 2013).

Similar to studies of SA and SI, findings from the literature examining suicide risk and the LDAEP are mixed. One study among a sample of college undergraduates and community members did not find a relationship between the LDAEP and Suicidal Behaviors Questionnaire scores (*g* = 0.14; Marsic, 2012; Osman et al., 2001). A study among a patient sample also did find not any differences in the LDAEP between those of varying levels of suicide risk (*g* = - 0.23; Graßnickel et al., 2015). In contrast, one study among a sample of male undergraduates found a positive relationship between the LDAEP and Suicidal Behaviors Questionnaire scores (*g* = 0.14; Marsic et al., 2015).

#### Late Positive Potential (LPP)

As mentioned in our introduction, the LPP is an ERP that exhibits increased amplitudes to emotional stimuli, compared to neutral stimuli (Cuthbert et al., 2000; Weinberg & Hajcak, 2010, 2011). Specifically, the LPP is thought to reflect a protracted orienting response that begins with the P300 (see P300 section below), with both the LPP and P300 reflecting activity of both appetitive and aversive motivational systems (Hajcak & Foti, 2020). So far, only three studies have examined the relationship between the LPP and SI. In one study during which participants, who consisted of 40 undergraduate students, were instructed to reduce their emotional response to negatively valenced images, those with SI had a greater LPP in response to negatively valanced images, compared to students without SI (*g* = 0.86; Kudinova et al., 2016). However, the study did not find any differences between students with SI and those without SI for any other stimulus valence or instruction type (e.g., increased emotional response; passive viewing). Another, similar study among community adults presenting to a clinical trial found that the LPP was negatively associated with self-reported suicidal ideation on the Beck Suicide Scale but only in response to positively valanced stimuli during a condition during which participants were instructed to increase their emotional response (*g* = -0.26 – -0.28; Albanese et al., 2019b; Beck et al., 1988). In a separate, unpublished dataset of community adults, where participants either passively viewed threatening or neutral stimuli, or were told to downregulate their response to threatening stimuli, they did not find any relationship between any LPP and SI (Albanese & Schmidt, 2020).

Regarding SAs, only one study has examined differences in the LPP between those with a SA history and those without a SA history. Among a sample of psychiatric outpatients, Weinberg and colleagues (2017) found that the LPP in response to threat was significantly blunted among patients with a previous SA compared to patients with SI who had no history of SA (*g* = -0.48); furthermore, there were no differences in LPP amplitude to positive stimuli between the groups. Finally, two studies have examined the relationship between suicide risk and the LPP. One study in a sample of psychiatric outpatients found that the LPP to positive and negative stimuli both had a negative relationship with suicide risk (*g* = -0.43 – -0.45; Weinberg et al., 2016). Finally, in the same unpublished study we discussed while reviewing SI, they found no relationship between the LPP in response to any stimulus type or condition and suicide risk level (Albanese & Schmidt, 2020).

#### P300

The P300 is a positive deflection in the ERP waveform that is classically elicited to the presentation of infrequent stimuli during an oddball task (Donchin, 1981). Some evidence suggests that the P300 and LPP may both index salience or significance and thus may reflect similar underlying neural processes (Bradley, 2009; Hajcak & Foti, 2020). Three studies have examined the P300 and how it may relate to SI. The first did not find any relationship between the P300 and self-reported SI (*g* = -0.57; Baik et al., 2018). Another study among psychiatric patients found a significant, positive relationship between the P300 to a variety of stimuli during an affective incentive delay task and SI (Song et al., 2019). Specifically, the study found that SI was positively related to the P300 to the target stimulus, to the positive feedback P300 regardless of condition, and to the negative feedback P300 during a reward condition (i.e., negative feedback = a neutral picture; positive feedback = rewarding picture). Finally, an unpublished study among community adults found a positive relationship between the P300 to viewing pictures with threatening stimuli and SI (Albanese & Schmidt, 2020).

For SA history, two early studies in small samples found a reduced auditory P300 in patients with a previous SA, compared to patients without a previous SA (*g* = -1.97 – -1.35; Hansenne et al., 1994, 1996). In contrast, another study found that the auditory P300 was greater in those with a previous SA, compared to those without a previous SA, but only at Fz (*g* = 0.83; Chen et al., 2005). In line with this study, another study found an unrealistically large (viz., Hedges’ *g* = 3.57) difference in the P300 during an oddball task between those with a previous SA and those without a previous SA (Jandl et al., 2010). Two more recent studies have also reported conflicting results. Albanese and colleagues (2019a) did not find a difference in the P300 between participants with SA and those with SI but no history of SA (*g* = -0.27 – 0.20). In contrast, a separate study comparing those with SA to controls found that those with SA had a blunted cue P300 during a monetary incentive delay task (*g* = -0.54; Tsypes et al., 2020).

In regard to suicide risk, a study in a sample of adolescent inpatients used an auditory oddball paradigm and found that those with greater suicide risk (i.e., adolescents with recent SI and/or SA) had greater P300 than controls (*g* = 1.26; Tavakoli et al., 2018). A similar study among adolescent inpatients using a visual oddball task did not find any differences in the P300 (*g* = 0.13 – 0.27; Tavakoli et al., 2020). Last, in an unpublished study, Albanese (2020) found that most P300s did not correlate with suicide risk measure scores, with one exception: The study found a negative correlation (i.e., *r* = -0.23) between the P300 to neutral stimuli in a visual task and a self-report measure of suicide risk.

#### Reward Positivity (RewP)

The RewP is a relative positive deflection in the ERP that is maximal approximately 300 ms after presentation of rewards compared to nonrewards (Holroyd et al., 2008; Proudfit, 2015). A more positive RewP is related to increased BOLD response in the ventral striatum (Becker et al., 2014; Carlson et al., 2011). A blunted RewP has been inconsistently related to SI. In a sample of children with SI and matched controls, the RewP was smaller in children with recent SI (i.e., SI within the last two weeks), compared to controls (*g* = -0.60; Tsypes et al., 2019). However, in a conceptual replication using two larger samples of slightly older children, there was no difference in the RewP between children with recent SI and controls (*g* = -0.27 – 0.16; Gallyer, Burani, et al., 2020). Similarly, a study using pictures as the rewarding or unrewarding stimuli in a sample of patients with major depressive disorder did not find a relationship between the RewP and SI (*g* = -0.18 – -0.16; Song et al., 2019).

Researchers have only recently begun to examine the RewP and its potential relationship to SA and suicide risk. One study in a community adult sample found no difference in the RewP between those with a previous SA and controls without a previous SA (*g* = -0.14 – 0.21; Tsypes et al., 2020). An unpublished study with a similar design by the same group also failed to find a difference in the RewP between those with a SA and controls (*g* = -0.49 – -0.02; Tsypes & Gibb, 2020). Last, among a sample of adolescents experiencing depression, there was a positive relationship between the RewP and having suicide risk (i.e., experiencing any STB; *g* = 0.74; Pegg et al., 2020).

#### Contingent Negative Variation (CNV), Post-Imperative Negative Variation (PINV), and Stimulus-Preceding Negativity (SPN)

The CNV is a negative-going wave that precedes the presentation of an affective cue or action initiation, with the CNV demonstrating greater (i.e., more negative) amplitudes during the anticipation of more emotionally salient stimuli (Angus et al., 2017; Baas et al., 2002; Tecce, 1972). Work has shown that, in some circumstances, the CNV can continue for an extended time period, resulting in what has been called the PINV (Timsit et al., 1970). Research using simultaneous EEG and fMRI found that trial-by-trial variation of the CNV is associated with BOLD activity in the thalamus, anterior cingulate, and supplementary motor cortex (Nagai et al., 2004). There are few studies that have examined the CNV in STBs. The single study that examined the CNV in SI found no relationship between the CNV and SI, regardless of the condition, during an affective incentive delay task (*g* = -0.39 – -0.04; Song et al., 2019). A series of early studies with very small sample sizes consistently found that the CNV was blunted in patients (i.e., closer to zero) with a previous SA, compared to patients without a SA (*g* = -0.59 – 1.27; Ashton et al., 1994; Hansenne et al., 1994, 1996). Notably, one of these studies also examined the PINV but did not find any differences (*g* = 0.46; Ashton et al., 1994). A more recent study, however, failed to find any differences in the CNV between SA groups, and there have yet to be any studies examining suicide risk and the CNV (*g* = 0.20; Tsypes et al., 2020). In relation to the SPN (i.e., a slow-wave ERP that precedes anticipation in several different tasks; van Boxtel & Böcker, 2004), the single study to examine differences in this ERP between those with STBs found no differences in the SPN between participants with a previous SA, compared to those without a previous SA (*g* = 0.00; Tsypes et al., 2020).

#### N200, P200, and N100

The N200 is a negative-deflection ERP that reflects conflict-monitoring processing, with the N200 exhibiting more negative amplitudes to no-go cues, relative to go trials, during go/no-go tasks (Donkers & Boxtel, 2004; Nieuwenhuis et al., 2003). Only two studies have examined the N200 and its relation to STBs. The first found that the N200 during a go/no-go task was reduced (i.e., less negative) among participants with a previous SA, compared to participants experiencing SI without a previous SA (*g* = 0.66; Albanese et al., 2019a). The second study did not find any evidence for differences in the N200 between patients hospitalized for acute suicide risk and matched healthy controls (*g* = -0.60; Tavakoli et al., 2020). A single study has examined the N100 (an early sensory ERP; Pratt, 2011) and STBs. This study failed to find differences in the N100 in a small sample of patients with a previous SA versus controls (*g* = 0.27; Hansenne et al., 1996).

The P200 is a positive-deflection ERP shortly following stimulus presentation that is maximal approximately 180 ms after stimulus onset (Carretié et al., 2006). Research has shown that the P200 has the greatest amplitude to novel target stimuli (Luck & Hillyard, 1994). Research has also shown that the P200 exhibits greater amplitudes to affective stimuli, relative to neutral stimuli (Carretié et al., 2001; Delplanque et al., 2004; Olofsson & Polich, 2007). As with the N200, very few studies have examined the P200 and how it may relate to STBs. One study found a positive correlation between SI and the P200 during an affective incentive delay task among a sample of psychiatric patients and healthy controls (*g* = 0.74; Song et al., 2019). Lastly, a study among a small sample of patients also found that the P200 was reduced in patients with a previous SA, compared to controls (*g* = -2.37; Hansenne et al., 1996).

### Summary of Meta-Analytic Results and Publication Bias

For forest plots of all ERP-STB meta-analysis results, see Figure 2 (SI), Figure 3 (SA), Figure 4 (suicide risk) and Figure 5 (SA vs. SI-only). Across all ERP-STB combinations there were no statistically significant effects (Table 2) with one exception: there was a significant blunting of the RewP in those with SI compared to those without SI (*g* = -0.06, 95% CI [-0.12, - 0.01], *I*^2^ = 4.29%; Figure 2D). Similarly, most of our modified Egger’s regression analyses to investigate small-sample publication bias were not significant, with one exception: in our meta-analysis testing for differences in the P300 between those with SA vs. those with SI-only, there was evidence of small sample publication bias (*F*[1, 1] = 931.02, *p* = .021), with studies with larger samples exhibiting smaller effect sizes (*b* = -0.66).

**Figure 2.**
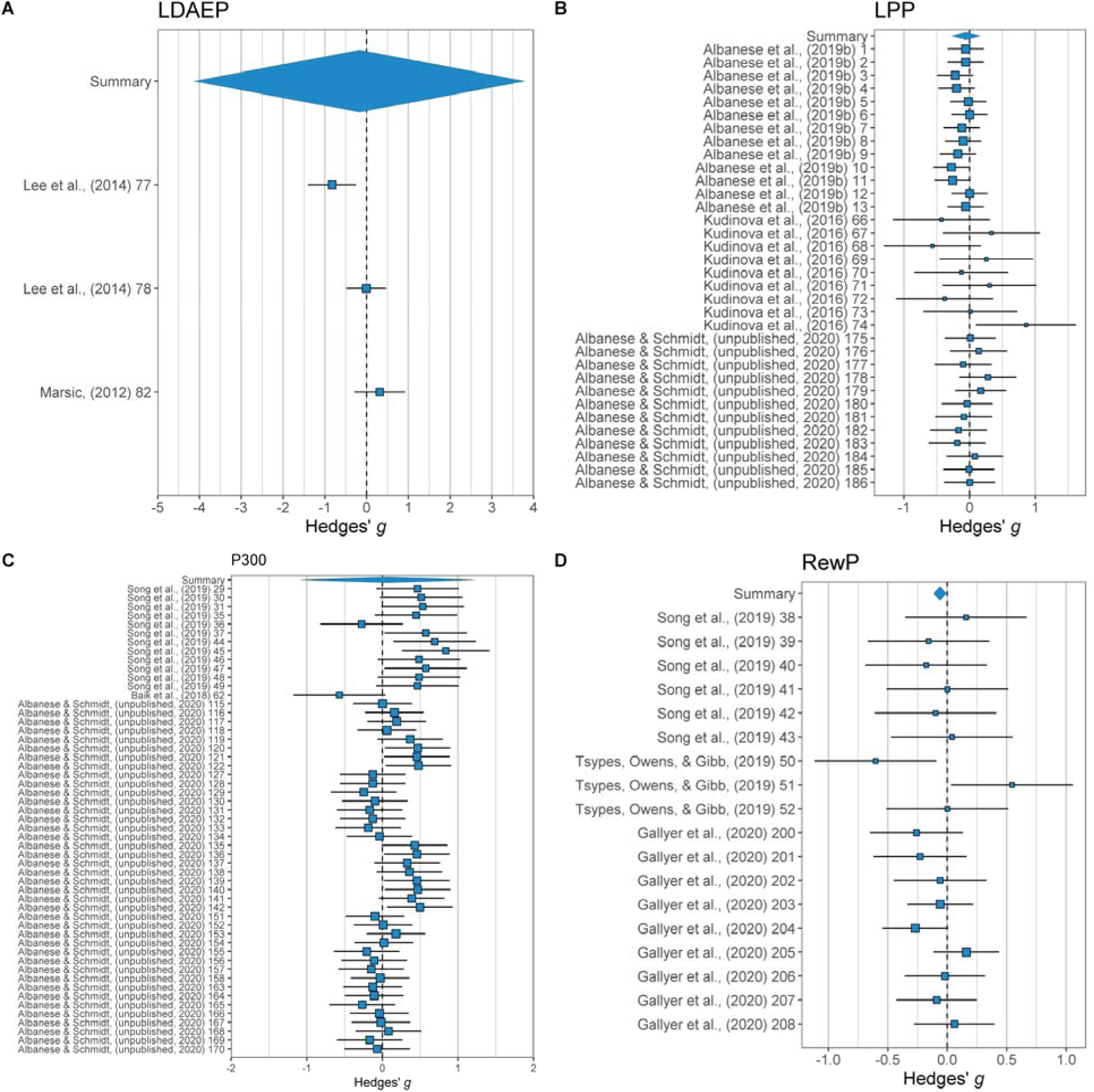
SI Meta-Analysis Forest Plots Across ERPs

**Figure 3.**
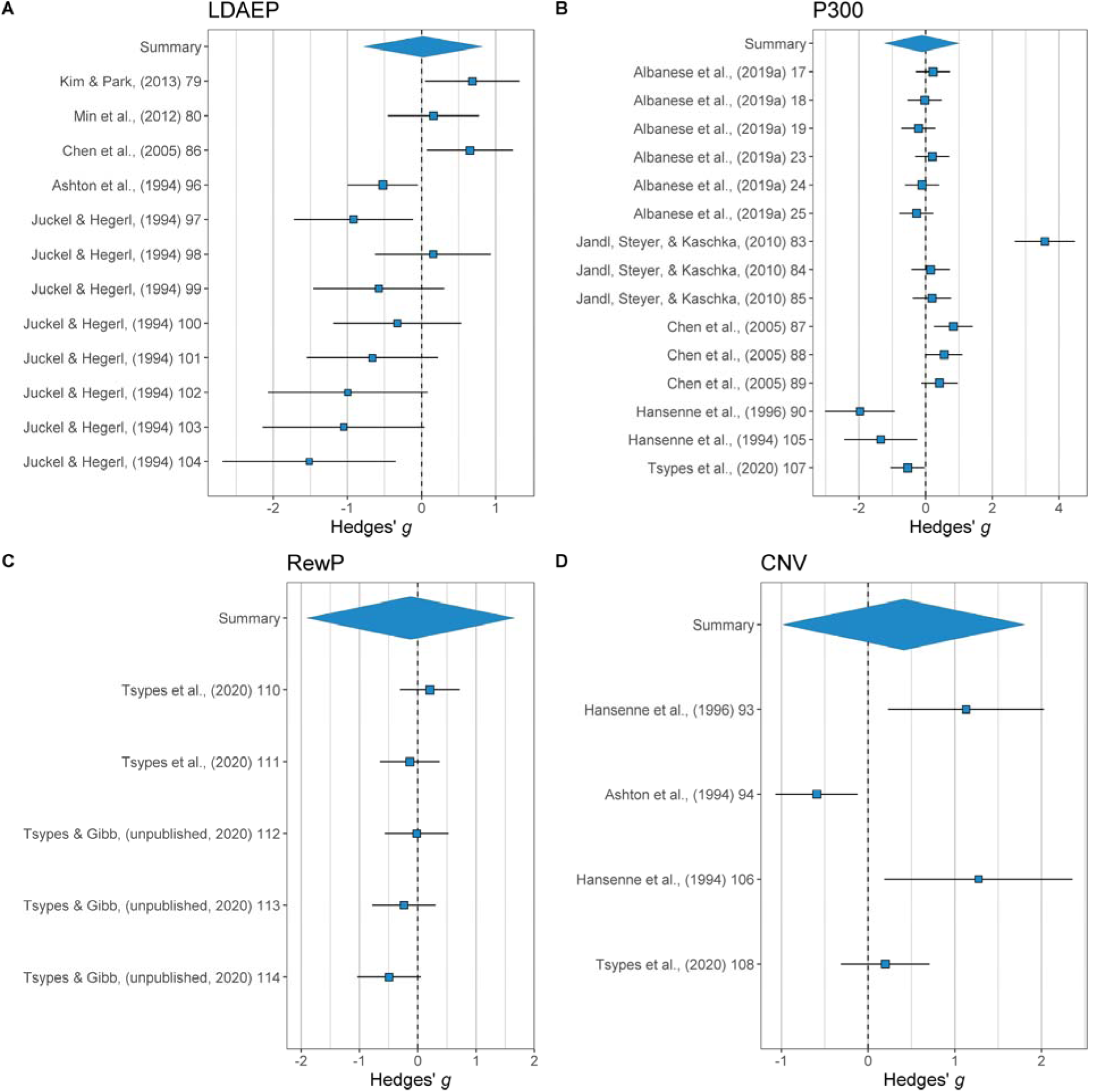
SA Meta-Analysis Forest Plots Across ERPs

**Figure 4.**
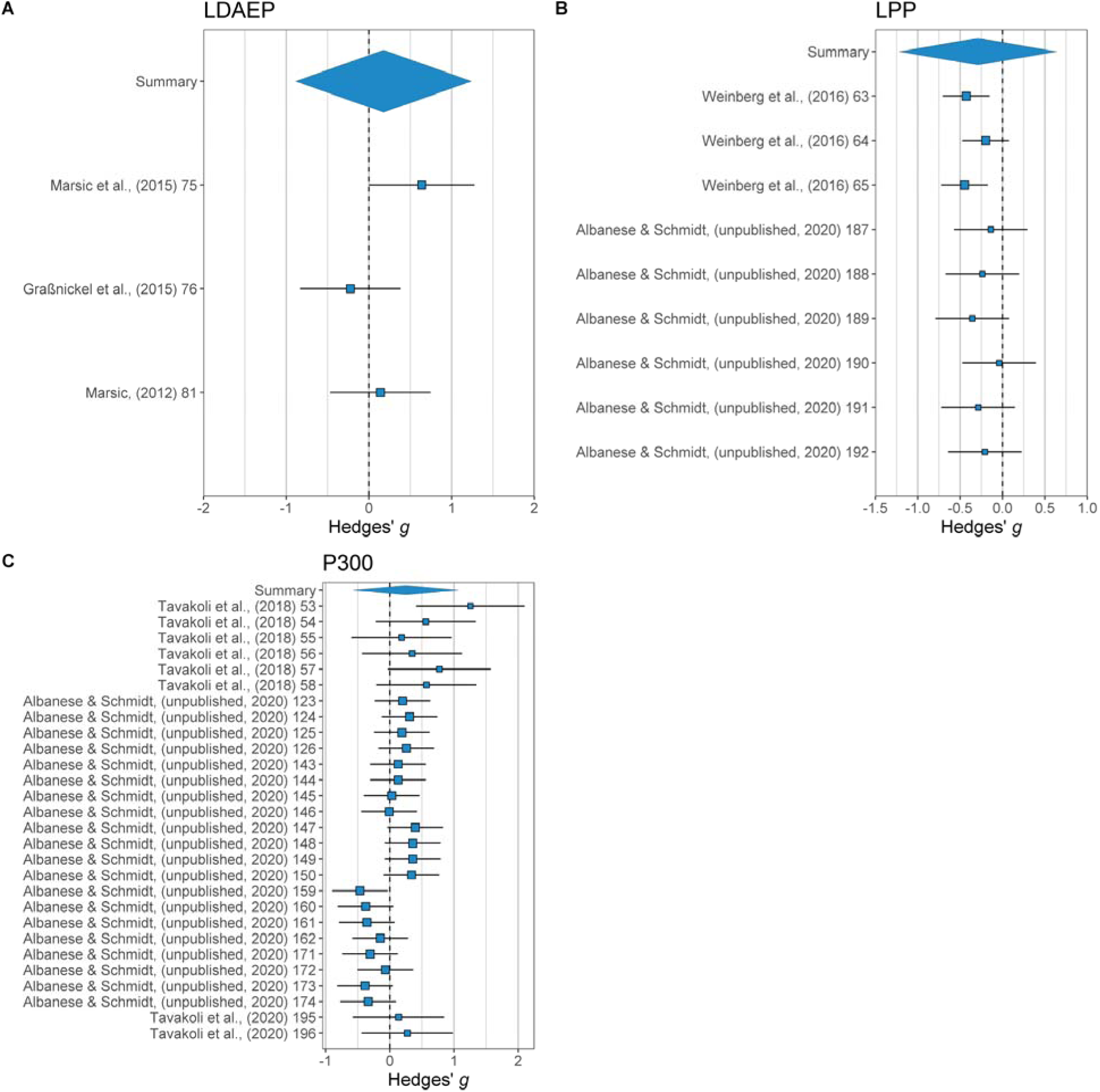
Suicide Risk Meta-Analysis Forest Plots Across ERPs

**Figure 5.**
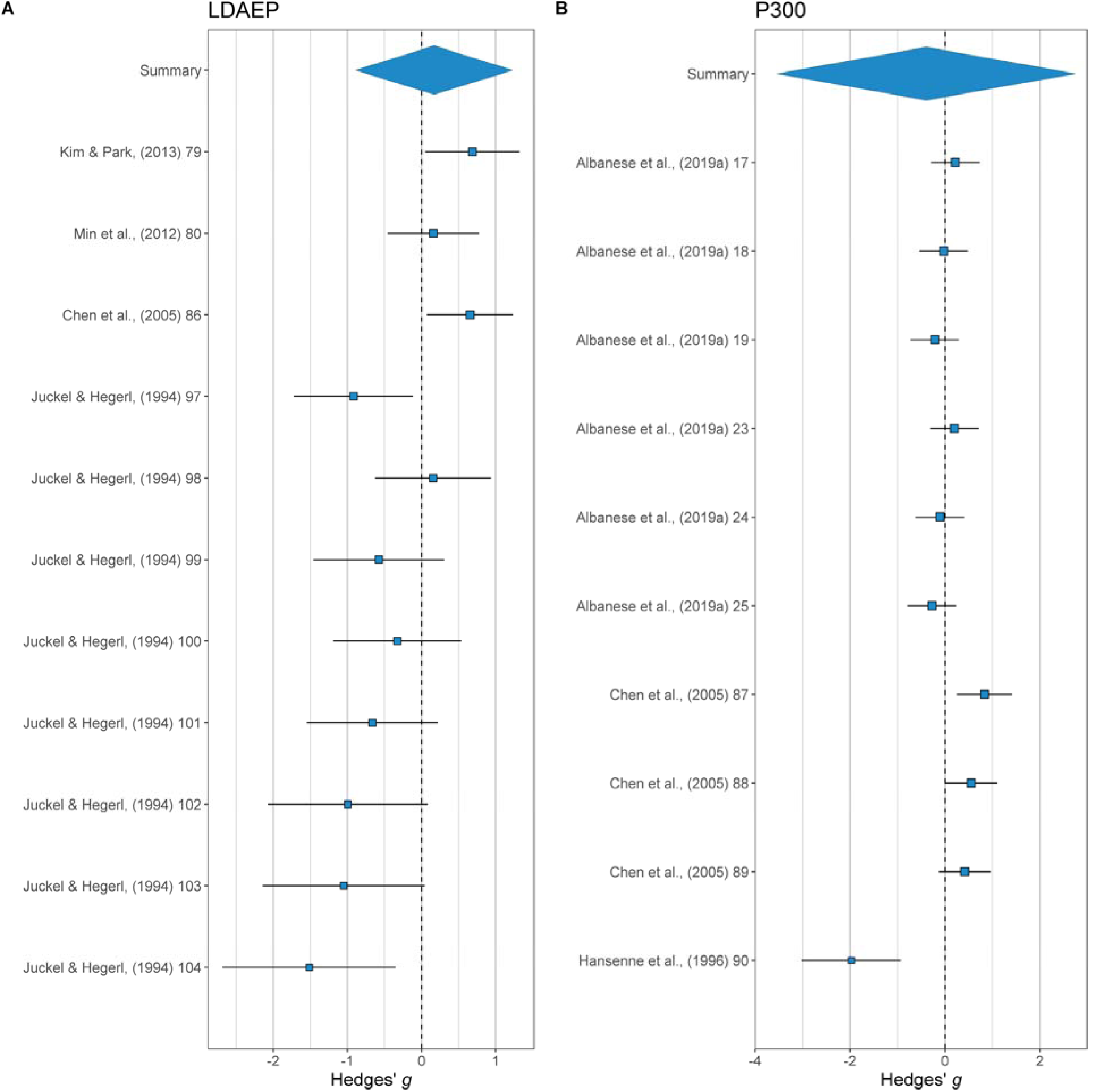
SA vs. SI-Only Meta-Analysis Forest Plots Across ERPs

**Table 2.**
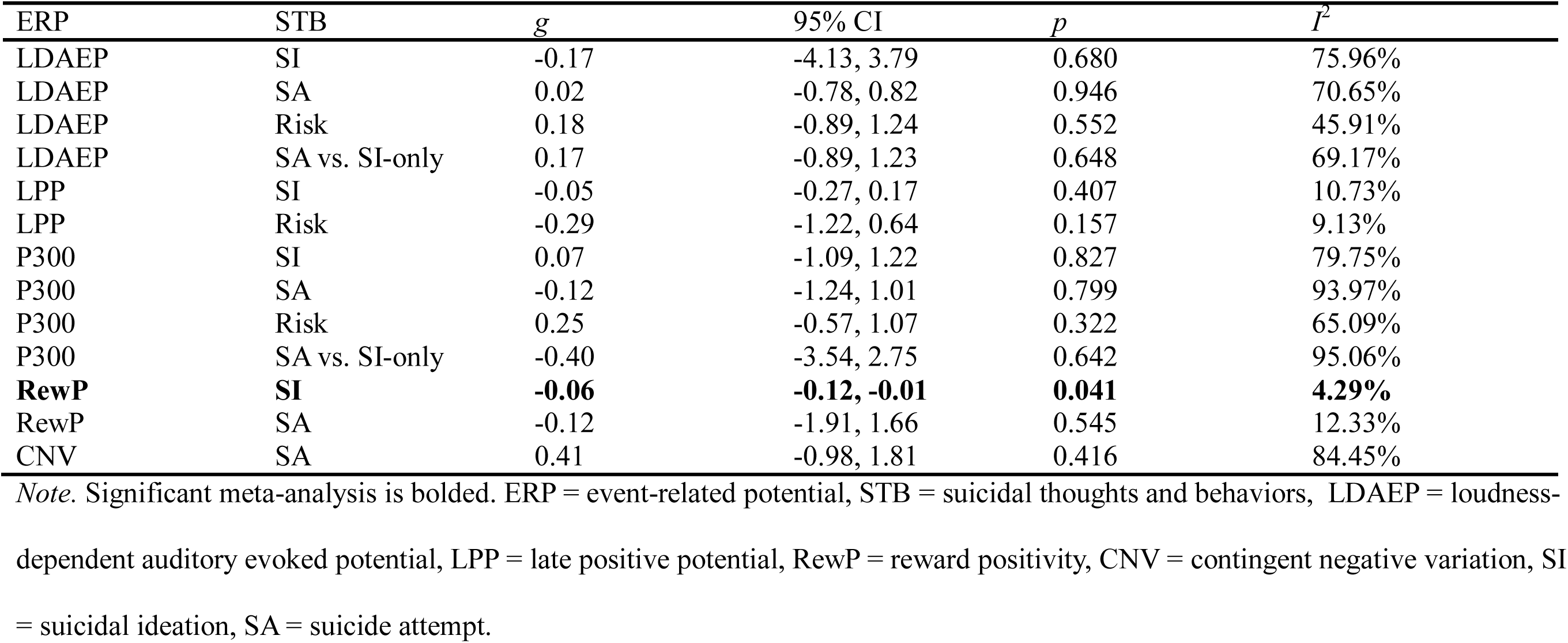
Meta-Analysis Results

### Statistical Power

The statistical power across a range of true effect sizes of the median sample size within each study is presented in Figure 6. This shows that most of the studies in this area are severely underpowered to detect effects that are probable in this area. For example, in the maximum true effect size tested (i.e., δ = 1.0), six studies had statistical power less than 0.80, the traditionally accepted cutoff for sufficient statistical power. Of the remaining twenty-one studies, the median true effect size needed to reach sufficient statistical power was δ = 0.40, seventeen of which did not reach sufficient statistical power until δ >0.60.

**Figure 6.**
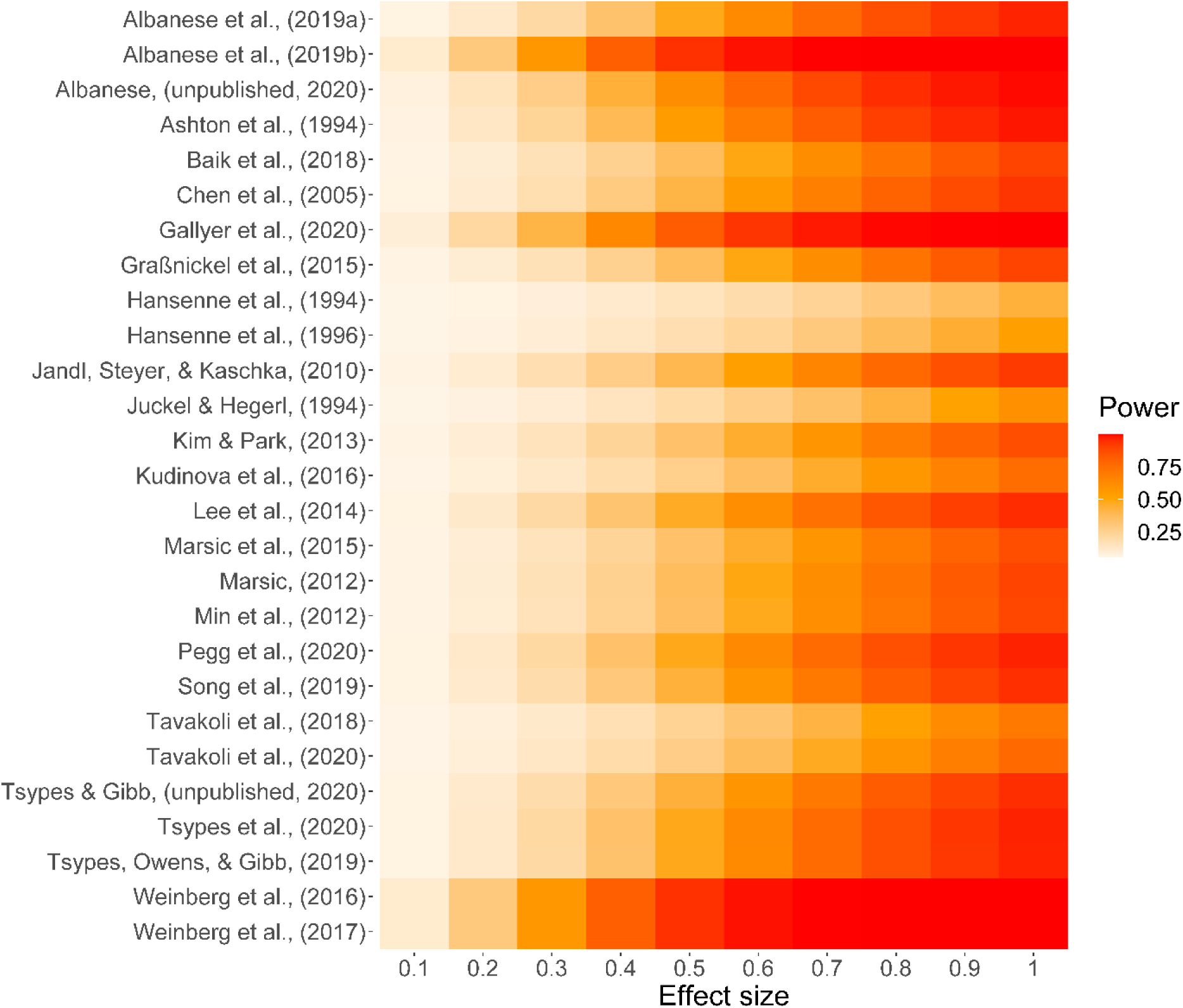
Median Power of Studies in Meta-Analysis Across True Effect Sizes

## Discussion

Scientists have proposed that suicidal thoughts and behaviors (STBs) are at least in part the result of differences in neural architecture and functioning (Mann, 2003; Van Heeringen & Mann, 2014). Evidence from fMRI studies has led to divergent conclusions, with some papers concluding that there are neural differences between those with and without STBs while other papers suggest that there is not yet enough evidence to conclude that there are differences in neural functioning between individuals with and without STBs (Huang et al., 2020; Schmaal et al., 2019). Event-related potentials (ERPs) offer an alternative approach to quantifying neural functioning in those with STBs that is more cost-effective, provides excellent temporal resolution, and directly measures neurophysiological processes. However, the effectiveness of ERPs at differentiating individuals with versus without STBs was unknown.

Our qualitative review of specific ERPs and their relations to ERPs revealed that, across ERPs and STB combinations, the literature is highly mixed. Our meta-analysis results corroborated this observation. Specifically, across meta-analyses we largely found null results, with one exception: there was a significant blunting of the RewP in those with SI compared to those without SI. This in line with a recent meta-analysis that found that the RewP was blunted in those with depression (Keren et al., 2018). Given the high correlation between SI and depression, this result is unsurprising, but this also means that it cannot be ruled out that differences in depression, rather than SI, are driving these results (Rogers et al., 2016). Together then, our results provided no strong evidence that any ERP assessed significantly differs across any STB group. We next discuss the possible reasons for our largely null results, their implications, and our recommendations for future research.

One problem may be that studies use vastly different behavioral tasks to elicit purportedly similar ERPs. Though doing so may be good in theory for generalizability purposes, doing so may introduce validity issues if similar ERPs from different tasks are elicited by slightly different neural processes. For example, is the RewP to the monetary reward task the same as a task during which the reward is a pleasant affective image? Data are lacking on this question and on many other task comparisons we reviewed. However, some evidence suggests that the ostensibly same ERP elicited from different tasks are only modestly correlated with one another (i.e., *r* = 0.33–0.43; Riesel et al., 2013). Thus, ERPs that are categorized the same in our meta-analysis may reflect distinct neural processes, thereby attenuating effects.

Another issue is that many studies used a single-item assessment or other problematic measurement approaches (e.g., combining measures without previous evidence of combined-measure validity) to measure STBs. Single-item assessments are particularly problematic, as participants may endorse SI or SA history but not meet standardized criteria for SI or SA when probed for more information (cf. Hom et al., 2016; Millner et al., 2015). Moreover, combining measures without sufficient justification and without establishing validity may attenuate effect sizes and make results less replicable (Flake & Fried, 2020). Moreover, many studies dichotomized continuous measures of STBs. On one hand, this practice is understandable, given the difficulty in collecting samples of individuals experiencing STBs relative to other populations within clinical psychology and psychiatry. With the sample sizes typically seen in these studies, there may simply not be many participants with varied scores on a STB measure. On the other hand, dichotomizing continuous variables may reduce statistical power and increase the false positive rate (Altman & Royston, 2006). Dichotomizing between presence versus absence of STBs may be particularly problematic, given evidence that suicide risk may be a rare instance of a construct that is categorical, with high suicide risk being categorically distinct from low suicide risk (Rufino et al., 2018; Witte et al., 2017).

Another explanation is that the literature is extremely underpowered. In support of this, we found that most studies were not sufficiently powered to detect realistic effect sizes, with some studies still demonstrating insufficient statistical power for unrealistically large (i.e., δ = 1.0) effect sizes. Given this poor statistical power, it is not surprising that the meta-analytic effect across nearly all our meta-analyses was not significant. Finally, our results may be explained by the possibility that there are truly no differences in ERPs across STB groups. Though we think there is currently insufficient evidence to draw this conclusion, it is a possibility that should be strongly considered given a recent well powered fMRI study that may point toward the same conclusion (Vidal-Ribas et al., 2021).

Before discussing the implications of our results, they should be interpreted in the context of our study’s limitations. First, because this area is relatively understudied, we were unable to conduct moderation analyses. This is an important area for future study, as it may emerge, for example, that particular ERPs do significantly differ in those with STBs, but only in certain subpopulations, like children or adolescents. Alternatively, specific ERPs may amplify the effects of suicide vulnerability factors on STBs in a diathesis-stress model. Second, as we will discuss in more depth below, null-hypothesis significance testing cannot provide evidence that there is no effect. Thus, while we largely found null results, it is possible that for some ERP-STB combinations, we were relatively underpowered to detect effects given the few studies in this area. Thus, our results do not definitively support the absence of effects. Third, one study provided an abnormally large number of effect sizes to our meta-analysis (i.e., Albanese & Schmidt, 2020). This study was an unpublished study that we had access to the data, and thus we were able to examine correlations between many STB measures and ERPs. However, because we accounted for our effect sizes being nested within studies in our analyses, we are confident that this study did not drive our null results.

### Future of the Study of STBs Using ERPs

Our results imply that this small-but-growing research area may need to consider significant reforms, and to adopt approaches that can better determine whether there are significant differences in ERPs in those with STBs, or, critically, whether there are *not* differences. One way to do this would be to use equivalence tests (Lakens, 2017). Though it is a common misunderstanding, traditional null-hypothesis significance testing does not allow conclusions about whether there is not an effect. Equivalence tests can provide evidence whether the effect is effectively zero by testing whether the observed effect is smaller than what the researcher determines is the smallest effect size of interest. Another approach would be to use Bayesian analyses. These analyses allow researchers to account for their prior beliefs about what the effect size is, and then receive results that imply whether there is more evidence for there being an effect or not.

Similar to the fMRI literature, in which a meta-analysis found a median sample size of 45 (Huang et al., 2020), we found that the majority of studies using ERPs to study STBs are severely underpowered. That is, most studies to date are too underpowered to provide meaningful information about the neurobiology of STBs. This is because low statistical power increases the frequency of both false negatives and false positives, which limits the replicability of studies of STBs (Button et al., 2013). There are likely many factors contributing to low statistical power in the neuroimaging, ERP, and STB literatures, including institutional incentives (Smaldino & McElreath, 2016), the high financial cost to conduct neuroimaging studies, and the relatively low base rate of STBs (Fazel & Runeson, 2020). To combat low statistical power in the field of neuroimaging, consortia have been developed to construct large neuroimaging datasets (e.g., Thompson et al., 2014). We believe that a similar consortium focused on using ERPs to study STBs, as well as greater collaboration in general, will be necessary to determine whether neurobiological differences play a role in the etiology of STBs.

However, larger datasets alone will not be sufficient to make the research in this area meaningful. We also encountered poor measurement practices, with most studies using single-item assessments. To combat this issue, researchers should prioritize using multi-item assessments and should, ideally, use interview-based measures that allow the researcher to ask clarifying questions (e.g., Chu et al., 2015; Fox et al., 2020; Gallyer, Chu, et al., 2020; Joiner et al., 1999; Nock et al., 2007). Though research in military samples suggests that suicide risk measures are inaccurate in making absolute determinations of who is at risk, this work has also suggested that some of the most popular measures of suicide risk are equally effective (Gutierrez et al., 2020). Thus, researchers might focus on using any multi-item measure, rather than exerting too much effort determining which is the best or combining different measures.

We also found that ERPs were elicited using very different tasks that may not reflect the same neural processes. Researchers may consider creating a standardized battery of tasks that are shown to produce ERPs with high internal consistency. This recommendation, in combination with our call for greater collaboration and/or a consortium dedicated to studying STBs using ERPs, would be particularly beneficial for the field. Doing so would make results across studies more comparable, and maximizing internal consistency would increase the ability to relate ERPs to individual differences such as STBs (Hajcak et al., 2017). By improving statistical power, measurement practices, and the consistency of behavioral tasks, we are confident that ERPs will have much to contribute to the understanding of whether neurobiology plays a role in STBs.

## References

Albanese, B. J., Macatee, R. J., Gallyer, A. J., Stanley, I. H., Joiner, T. E., & Schmidt, N. B. (2019). Impaired Conflict Detection Differentiates Suicide Attempters From Ideating Nonattempters: Evidence From Event-Related Potentials. Biological Psychiatry: Cognitive Neuroscience and Neuroimaging, 4(10), 902–912. https://doi.org/Suicidal behaviors among American Indian/Alaska Native firefighters: Evidence for the role of painful and provocative events

Albanese, B. J., Macatee, R. J., Stanley, I. H., Bauer, B. W., Capron, D. W., Bernat, E., Joiner, T. E., & Schmidt, N. B. (2019). Differentiating suicide attempts and suicidal ideation using neural markers of emotion regulation. Journal of Affective Disorders, 257, 536–550. https://doi.org/10.1016/j.jad.2019.07.014

Albanese, B. J., & Schmidt, N. B. (2020). *Unpublished Data*.

Altman, D. G., & Royston, P. (2006). The cost of dichotomising continuous variables. BMJ, 332, 1080.

Angus, D. J., Latham, A. J., Harmon-Jones, E., Deliano, M., Balleine, B., & Braddon-Mitchell, D. (2017). Electrocortical components of anticipation and consumption in a monetary incentive delay task. Psychophysiology, 54(11), 1686–1705. https://doi.org/10.1111/psyp.12913

Arnold, J. B. (2019). ggthemes: Extra Themes, Scales and Geoms for “ggplot2*.”* https://CRAN.R-project.org/package=ggthemes

Ashton, C. H., Marshall, E. F., Hassanyeh, F., Marsh, V. R., & Wright-Honari, S. (1994). Biological correlates of deliberate self-harm behaviour: A study of electroencephalographic, biochemical and psychological variables in parasuicide. Acta Psychiatrica Scandinavica, 90(5), 316–323. https://doi.org/10.1111/j.1600-0447.1994.tb01600.x

Baas, J. M. P., Kenemans, J. L., Böcker, K. B. E., & Verbaten, M. N. (2002). Threat-induced cortical processing and startle potentiation: Neuroreport, 13(1), 133–137. https://doi.org/10.1097/00001756-200201210-00031

Baik, S. Y., Jeong, M., Kim, H. S., & Lee, S.-H. (2018). ERP investigation of attentional disengagement from suicide-relevant information in patients with major depressive disorder. Journal of Affective Disorders, 225, 357–364. https://doi.org/10.1016/j.jad.2017.08.046

Bauer, B. W., Albanese, B. J., Macatee, R. J., Tucker, R. P., Bernat, E., Schmidt, N. B., & Capron, D. W. (2020). Fearlessness About Death is Related to Diminished Late Positive Potential Responses When Viewing Threatening and Mutilation Images in Suicidal Ideators. Cognitive Therapy and Research. https://doi.org/10.1007/s10608-020-10094-4

Beck, A. T., Steer, R. A., & Ranieri, W. F. (1988). Scale for suicide ideation: Psychometric properties of a self□report version. Journal of Clinical Psychology, 44(4), 7.

Becker, M. P. I., Nitsch, A. M., Miltner, W. H. R., & Straube, T. (2014). A Single-Trial Estimation of the Feedback-Related Negativity and Its Relation to BOLD Responses in a Time-Estimation Task. Journal of Neuroscience, 34(8), 3005–3012. https://doi.org/10.1523/JNEUROSCI.3684-13.2014

Bradley, M. M. (2009). Natural selective attention: Orienting and emotion. Psychophysiology, 46(1), 1–11. https://doi.org/10.1111/j.1469-8986.2008.00702.x

Button, K. S., Ioannidis, J. P. A., Mokrysz, C., Nosek, B. A., Flint, J., Robinson, E. S. J., & Munafò, M. R. (2013). Power failure: Why small sample size undermines the reliability of neuroscience. Nature Reviews Neuroscience, 14. https://doi.org/10.1038/nrn3475

Carlson, J. M., Foti, D., Mujica-Parodi, L. R., Harmon-Jones, E., & Hajcak, G. (2011). Ventral striatal and medial prefrontal BOLD activation is correlated with reward-related electrocortical activity: A combined ERP and fMRI study. NeuroImage, 57(4), 1608–1616. https://doi.org/10.1016/j.neuroimage.2011.05.037

Carretié, L., Hinojosa, J. A., Albert, J., & Mercado, F. (2006). Neural response to sustained affective visual stimulation using an indirect task. Experimental Brain Research, 174(4), 630–637. https://doi.org/10.1007/s00221-006-0510-y

Carretié, L., Mercado, F., Tapia, M., & Hinojosa, J. A. (2001). Emotion, attention, and the ‘negativity bias’, studied through event-related potentials. International Journal of Psychophysiology, 41(1), 75–85. https://doi.org/10.1016/S0167-8760(00)00195-1

Centers for Disease Control. (2017). WISQARS: Web-Based Injury Statistics Query and Reporting System [Government Database]. www.cdc.gov/injury/wisqars

Chang, B. P., Franklin, J. C., Ribeiro, J. D., Fox, K. R., Bentley, K. H., & Nock, M. K. (2016). Biological risk factors for suicidal behaviors: A meta-analysis. Translational Psychiatry, 6. https://doi.org/10.1038/tp.2016.165

Chen, T.-J., Yu, Y. W.-Y., Chen, M.-C., Wang, S.-Y., Tsai, S.-J., & Lee, T.-W. (2005). Serotonin Dysfunction and Suicide Attempts in Major Depressives: An Auditory Event-Related Potential Study. Neuropsychobiology, 52(1), 28–36. https://doi.org/10.1159/000086175

Chu, C., Klein, K. M., Buchman-Schmitt, J. M., Hom, M. A., Hagan, C. R., & Joiner, T. E. (2015). Routinized assessment of suicide risk in clinical practice: An empirically informed update. Journal of Clinical Psychology, 71(12), 1186–1200. https://doi.org/10.1002/jclp.22210

Clayson, P. E., Carbine, K. A., Baldwin, S. A., & Larson, M. J. (2019). Methodological reporting behavior, sample sizes, and statistical power in studies of event □related potentials: Barriers to reproducibility and replicability. Psychophysiology. https://doi.org/10.1111/psyp.13437

Cuthbert, B. N., Schupp, H. T., Bradley, M. M., Birbaumer, N., & Lang, P. J. (2000). Brain potentials in affective picture processing: Covariation with autonomic arousal and affective report. Biological Psychology, 52, 95–111.

Del Re, A. C. (2013). compute.es: Compute effect sizes. http://cran.r-project.org/web/packages/compute.es

Delplanque, S., Lavoie, M. E., Hot, P., Silvert, L., & Sequeira, H. (2004). Modulation of cognitive processing by emotional valence studied through event-related potentials in humans. Neuroscience Letters, 356(1), 1–4. https://doi.org/10.1016/j.neulet.2003.10.014

Donchin, E. (1981). Surprise!… surprise? Psychophysiology, 18(5), 493–513.

Donkers, F. C. L., & Boxtel, G. J. M. V. (2004). The N2 in go/no-go tasks reflects conflict monitoring not response inhibitionq. Brain and Cognition, 56, 12.

Elliott, M. L., Knodt, A. R., Ireland, D., Morris, M. L., Poulton, R., Ramrakha, S., Sison, M. L., Moffitt, T. E., Caspi, A., & Hariri, A. R. (2020). What Is the Test-Retest Reliability of Common Task-Functional MRI Measures? New Empirical Evidence and a Meta-Analysis. Psychological Science, 31(7), 792–806. https://doi.org/10.1177/0956797620916786

Fazel, S., & Runeson, B. (2020). Suicide. New England Journal of Medicine, 382(3), 266–274. https://doi.org/10.1056/NEJMra1902944

Flake, J. K., & Fried, E. I. (2020). Measurement Schmeasurement: Questionable Measurement Practices and How to Avoid Them. Advances in Methods and Practices in Psychological Science, 3(4), 10.

Fox, K. R., Harris, J. A., Wang, S. B., Millner, A. J., Deming, C. A., & Nock, M. K. (2020). Self-Injurious Thoughts and Behaviors Interview—Revised: Development, reliability, and validity. Psychological Assessment. https://doi.org/10.1037/pas0000819

Franklin, J. C., Ribeiro, J. D., Fox, K. R., Bentley, K. H., Kleiman, E. M., Huang, X., Musacchio, K. M., Jaroszewski, A. C., Chang, B. P., & Nock, M. K. (2017). Risk factors for suicidal thoughts and behaviors: A meta-analysis of 50 years of research. Psychological Bulletin, 143(2), 187–232. https://doi.org/10.1037/bul0000084

Gallyer, A. J., Burani, K., Mulligan, E. M., Santopetro, N., Dougherty, S. P., Jeon, M. E., Nelson, B. D., Joiner, T. E., & Hajcak, G. (2020). Examining Blunted Initial Response to Reward and Recent Suicidal Ideation in Children and Adolescents Using Event-Related Potentials: Failure to Conceptually Replicate Across Two Independent Samples. BioRxiv. https://doi.org/10.1101/2020.05.19.104208

Gallyer, A. J., Chu, C., Klein, K. M., Quintana, J., Carlton, C., Dougherty, S. P., & Joiner, T. E. (2020). Routinized categorization of suicide risk into actionable strata: Establishing the validity of an existing suicide risk assessment framework in an outpatient sample. Journal of Clinical Psychology, jclp.22994. https://doi.org/10.1002/jclp.22994

Gallyer, A. J., Hajcak, G., & Joiner, T. E. (2020). What is capability for suicide? A review of the current evidence. PsyArXiv. https://doi.org/10.31234/osf.io/xgwa5

Gibb, B. E., & Tsypes, A. (2019). Using Event-Related Potentials to Improve Our Prediction of Suicide Risk. Biological Psychiatry: Cognitive Neuroscience and Neuroimaging, 4(10), 854–855. https://doi.org/10.1016/j.bpsc.2019.08.003

Graßnickel, V., Illes, F., Juckel, G., & Uhl, I. (2015). Loudness dependence of auditory evoked potentials (LDAEP) in clinical monitoring of suicidal patients with major depression in comparison with non-suicidal depressed patients and healthy volunteers: A follow-up-study. Journal of Affective Disorders, 184, 299–304. https://doi.org/10.1016/j.jad.2015.06.007

Gutierrez, P. M., Joiner, T., Hanson, J., Avery, K., Fender, A., Harrison, T., Kerns, K., McGowan, P., Stanley, I. H., Silva, C., & others. (2020). Clinical utility of suicide behavior and ideation measures: Implications for military suicide risk assessment. Psychological Assessment.

Hajcak, G., & Foti, D. (2020). Significance?… Significance! Empirical, methodological, and theoretical connections between the late positive potential and P300 as neural responses to stimulus significance: An integrative review. Psychophysiology, e13570. https://doi.org/10.1111/psyp.13570

Hajcak, G., Klawohn, J., & Meyer, A. (2019). The Utility of Event-Related Potentials in Clinical Psychology. Annual Review of Clinical Psychology, 15(1), 71–95. https://doi.org/10.1146/annurev-clinpsy-050718-095457

Hajcak, G., Meyer, A., & Kotov, R. (2017). Psychometrics and the neuroscience of individual differences: Internal consistency limits between-subjects effects. Journal of Abnormal Psychology, 126(6), 823–834. https://doi.org/10.1037/abn0000274

Hansenne, M., Pitchot, W., Gonzalez Moreno, A., Urcelay Zaldua, I., & Ansseau, M. (1996). Suicidal behavior in depressive disorder: An event-related potential study. Biological Psychiatry, 40(2), 116–122. https://doi.org/10.1016/0006-3223(95)00372-X

Hansenne, M., Pitchot, W., Moreno, A. G., Torrecilas, J. G., Mirel, J., & Ansseau, M. (1994). Psychophysiological correlates of suicidal behavior in depression. Neuropsychobiology, 30(1), 1–3.

Hedegaard, H., Curtin, S. C., & Warner, M. (2018). Suicide Mortality in the United States, 1999-2017 [Government Report]. Centers for Disease Control.

Hedges, L. V., Tipton, E., & Johnson, M. C. (2010). Robust variance estimation in meta-regression with dependent effect size estimates. Research Synthesis Methods, 1(1), 39–65. https://doi.org/10.1002/jrsm.5

Hegerl, U, Gallinat, J., & Juckel, G. (2001). Event-related potentials Do they reflect central serotonergic neurotransmission and do they predict clinical response to serotonin agonists? Journal of Affective Disorders, 8.

Hegerl, Ulrich, & Juckel, G. (1993). Intensity dependence of auditory evoked potentials as an indicator of central serotonergic neurotransmission: A new hypothesis. Biological Psychiatry, 15.

Holroyd, C. B., Pakzad-Vaezi, K. L., & Krigolson, O. E. (2008). The feedback correct-related positivity: Sensitivity of the event-related brain potential to unexpected positive feedback. Psychophysiology, 45(5), 688–697. https://doi.org/10.1111/j.1469-8986.2008.00668.x

Hom, M. A., Joiner, T. E., & Bernert, R. A. (2016). Limitations of a single-item assessment of suicide attempt history: Implications for standardized suicide risk assessment. Psychological Assessment, 28(8), 1026–1030. https://doi.org/10.1037/pas0000241

Huang, X., Rootes-Murdy, K., Bastidas, D. M., Nee, D. E., & Franklin, J. C. (2020). Brain Differences Associated with Self-Injurious Thoughts and Behaviors: A Meta-Analysis of Neuroimaging Studies. Scientific Reports, 10(1), 2404. https://doi.org/10.1038/s41598-020-59490-6

Jandl, M., Steyer, J., & Kaschka, W. P. (2010). Suicide risk markers in Major Depressive Disorder: A study of Electrodermal Activity and Event-Related Potentials. Journal of Affective Disorders, 123(1–3), 138–149. https://doi.org/10.1016/j.jad.2009.09.011

Joiner, T. E. (2005). Why people die by suicide. Harvard University Press.

Joiner, Thomas E, Brown, J. S., & Wingate, L. R. (2005). The psychology and neurobiology of suicidal behavior. Annual Review of Psychology, 56, 287–314. https://doi.org/10.1146/annurev.psych.56.091103.070320

Joiner, Thomas E., Walker, R. L., Rudd, M. D., & Jobes, D. A. (1999). Scientizing and routinizing the assessment of suicidality in outpatient practice. Professional Psychology: Research and Practice, 30(5), 447–453.

Juckel, G., & Hegerl, U. (1994). Evoked Potentials, Serotonin, and Suicidality. Pharmacopsychiatry, 27(S 1), 27–29. https://doi.org/10.1055/s-2007-1014323

Juckel, Georg, Molnar, M., Hegerl, U., Csepe, V., & Karmos, G. (1997). Auditory-Evoked Potentials as Indicator of Brain Serotonergic Activity First Evidence in Behaving Cats. Biological Psychiatry, 41, 15.

Keren, H., O’Callaghan, G., Vidal-Ribas, P., Buzzell, G. A., Brotman, M. A., Leibenluft, E., Pan, P. M., Meffert, L., Kaiser, A., Wolke, S., Pine, D. S., & Stringaris, A. (2018). Reward Processing in Depression: A Conceptual and Meta-Analytic Review Across fMRI and EEG Studies. American Journal of Psychiatry, 175(11), 1111–1120. https://doi.org/10.1176/appi.ajp.2018.17101124

Kim, D.-H., & Park, Y.-M. (2013). The association between suicidality and serotonergic dysfunction in depressed patients. Journal of Affective Disorders, 148(1), 72–76. https://doi.org/10.1016/j.jad.2012.11.051

Klonsky, E. D., & May, A. M. (2015). The Three-Step Theory (3ST): A new theory of suicide in the “ideation-to-action” framework. International Journal of Cognitive Therapy, 8(2), 114–129.

Klonsky, E. D., Saffer, B. Y., & Bryan, C. J. (2018). Ideation-to-action theories of suicide: A conceptual and empirical update. Current Opinion in Psychology, 22, 38–43. https://doi.org/10.1016/j.copsyc.2017.07.020

Kossmeier, M., Tran, U. S., & Voracek, M. (2020). metaviz: Forest Plots, Funnel Plots, and Visual Funnel Plot Inference for Meta-Analysis. https://CRAN.R-project.org/package=metaviz

Kudinova, A. Y., Owens, M., Burkhouse, K. L., Barretto, K. M., Bonanno, G. A., & Gibb, B. E. (2016). Differences in emotion modulation using cognitive reappraisal in individuals with and without suicidal ideation: An ERP study. Cognition and Emotion, 30(5), 999–1007. https://doi.org/10.1080/02699931.2015.1036841

Lakens, D. (2017). Equivalence Tests: A Practical Primer for *t* Tests, Correlations, and Meta-Analyses. Social Psychological and Personality Science, 8(4), 355–362. https://doi.org/10.1177/1948550617697177

Lee, S.-H., Park, Y.-C., Yoon, S., Kim, J.-I., & Hahn, S. W. (2014). Clinical implications of loudness dependence of auditory evoked potentials in patients with atypical depression. Progress in Neuro-Psychopharmacology and Biological Psychiatry, 54, 7–12. https://doi.org/10.1016/j.pnpbp.2014.05.010

Liu, Y., Huang, H., Mcginnis-Deweese, M., Keil, A., & Ding, M. (2012). Neural substrate of the late positive potential in emotional processing. Journal of Neuroscience, 32(42), 14563–14572. https://doi.org/10.1523/JNEUROSCI.3109-12.2012

Luck, S. J. (2014). An introduction to the event-related potential technique (2nd ed.). The MIT Press.

Luck, S. J., & Hillyard, S. A. (1994). Electrophysiological correlates of feature analysis during visual search. Psychophysiology, 31(3), 291–308.

Luck, S. J., & Kappenman, E. S. (2011). The Oxford Handbook of Event-Related Potential Components. Oxford University Press.

Mann, J. J. (2003). Neurobiology of suicidal behaviour. Nature Reviews: Neuroscience, 4, 819–828. https://doi.org/10.1038/nrn1220

Mann, J. J., & Rizk, M. M. (2020). A Brain-Centric Model of Suicidal Behavior. American Journal of Psychiatry, 15.

Marek, S., Tervo-Clemmens, B., Calabro, F. J., Montez, D. F., Kay, B. P., Hatoum, A. S., Donohue, M. R., Foran, W., Miller, R. L., Feczko, E., Miranda-Dominguez, O., Graham, A. M., Earl, E. A., Perrone, A. J., Cordova, M., Doyle, O., Moore, L. A., Conan, G., Uriarte, J., … Dosenbach, N. U. F. (2020). Towards Reproducible Brain-Wide Association Studies. BioRxiv. https://doi.org/10.1101/2020.08.21.257758

Marsic, A. (2012). THE RELATIONSHIP BETWEEN SUICIDE IDEATION AND PARASUICIDE: AN ELECTROPHYSIOLOGICAL INVESTIGATION USING THE LOUDNESS DEPENDENCE OF AUDITORY EVOKED POTENTIAL.

Marsic, A., Berman, M. E., Barry, T. D., & McCloskey, M. S. (2015). The Relationship Between Intentional Self-Injurious Behavior and the Loudness Dependence of Auditory Evoked Potential in Research Volunteers: Self-Injurious Behavior and the LDAEP. Journal of Clinical Psychology, 71(3), 250–257. https://doi.org/10.1002/jclp.22136

May, A. M., & Klonsky, E. D. (2016). What distinguishes suicide attempters from suicide Ideators? A meta-analysis of potential factors. Clinical Psychology: Science and Practice, 23(1), 5–20. https://doi.org/10.1111/cpsp.12136

Meyer, A., Riesel, A., & Proudfit, G. H. (2013). Reliability of the ERN across multiple tasks as a function of increasing errors: Reliability of the ERN across multiple tasks. Psychophysiology, 50(12), 1220–1225. https://doi.org/10.1111/psyp.12132

Millner, A. J., Lee, M. D., & Nock, M. K. (2015). Single-item measurement of suicidal behaviors: Validity and consequences of misclassification. PLoS ONE, 10(10). http://journals.plos.org/plosone/article/file?id=10.1371/journal.pone.0141606&type=printable

Min, J.-A., Lee, S.-H., Lee, S.-Y., Chae, J.-H., Lee, C.-U., Park, Y.-M., & Bae, S.-M. (2012). Clinical characteristics associated with different strengths of loudness dependence of auditory evoked potentials (LDAEP) in major depressive disorder. Psychiatry Research, 200(2–3), 374–381. https://doi.org/10.1016/j.psychres.2012.06.038

Moher, D., Liberati, A., Tetzlaff, J., & Altman, D. G. (2009). Preferred Reporting Items for Systematic Reviews and Meta-Analyses: The PRISMA Statement. Annals of Internal Medicine, 264–269.

Moran, T. P., Jendrusina, A. A., & Moser, J. S. (2013). The psychometric properties of the late positive potential during emotion processing and regulation. Brain Research, 1516, 66–75. https://doi.org/10.1016/j.brainres.2013.04.018

Nagai, Y., Critchley, H. D., Featherstone, E., Fenwick, P. B. C., Trimble, M. R., & Dolan, R. J. (2004). Brain activity relating to the contingent negative variation: An fMRI investigation. NeuroImage, 21(4), 1232–1241. https://doi.org/10.1016/j.neuroimage.2003.10.036

Nieuwenhuis, S., Yeung, N., Wildenberg, W. V. D., & Ridderinkhof, K. R. (2003). Electrophysiological correlates of anterior cingulate function in a go/no-go task: Effects of response conflict and trial type frequency. Cognitive, Affective, & Behavioral Neuroscience, 3(1), 10.

Nock, M. K., Holmberg, E. B., Photos, V. I., & Michel, B. D. (2007). Self-injurious thoughts and behaviors interview: Development, reliability and validity in an adolescent sample. Psychological Assessment, 19(3), 309–317. https://doi.org/10.1037/1040-3590.19.3.309

O’Connor, R. C., & Kirtley, O. J. (2018). The integrated motivational–volitional model of suicidal behaviour. Philosophical Transactions of the Royal Society B: Biological Sciences, 373(1754), 20170268. https://doi.org/10.1098/rstb.2017.0268

Olofsson, J. K., & Polich, J. (2007). Affective visual event-related potentials: Arousal, repetition, and time-on-task. Biological Psychology, 75(1), 101–108. https://doi.org/10.1016/j.biopsycho.2006.12.006

Osman, A., Bagge, C. L., Gutierrez, P. M., Konick, L. C., Kopper, B. A., & Barrios, F. X. (2001). The Suicidal Behaviors Questionnaire-Revised (SBQ-R): Validation with clinical and nonclinical samples. Assessment, 8(4), 443–454. https://doi.org/10.1177/107319110100800409

Pegg, S., Dickey, L., Green, H., & Kujawa, A. (2020). Differentiating clinically depressed adolescents with and without active suicidality: An examination of neurophysiological and self-report measures of reward responsiveness. Depression and Anxiety. https://doi.org/10.1002/da.23012

Pratt, H. (2011). Sensory ERP components. In The Oxford handbook of event-related potential components (pp. 89–114). Oxford University Press.

Proudfit, G. H. (2015). The reward positivity: From basic research on reward to a biomarker for depression: The reward positivity. Psychophysiology, 52(4), 449–459. https://doi.org/10.1111/psyp.12370

Pustejovsky, J. E., & Rodgers, M. A. (2019). Testing for funnel plot asymmetry of standardized mean differences. Research Synthesis Methods, 10(1), 57–71. https://doi.org/10.1002/jrsm.1332

Quintana, D. S. (2020a). metameta: A meta-analysis package for R (Version 0.1.1). 10.5281/zenodo.3944097

Quintana, D. S. (2020b). Most oxytocin administration studies are statistically underpowered to reliably detect (or reject) a wide range of effect sizes. Comprehensive Psychoneuroendocrinology, 4, 100014. https://doi.org/10.1016/j.cpnec.2020.100014

R Core Team. (2018). R: A language and environment for statistical computing. R Foundation for Statistical Computing. https://www.r-project.org/

Ribeiro, J. D., Huang, X., Fox, K. R., & Franklin, J. C. (2018). Depression and hopelessness as risk factors for suicide ideation, attempts and death: Meta-analysis of longitudinal studies. British Journal of Psychiatry. https://doi.org/10.1192/bjp.2018.27

Riesel, A., Weinberg, A., Endrass, T., Meyer, A., & Hajcak, G. (2013). The ERN is the ERN is the ERN? Convergent validity of error-related brain activity across different tasks. Biological Psychology, 93(3), 377–385. https://doi.org/10.1016/j.biopsycho.2013.04.007

Rodgers, M. A., & Pustejovsky, J. E. (2020). Evaluating meta-analytic methods to detect selective reporting in the presence of dependent effect sizes. Psychological Methods. https://doi.org/10.1037/met0000300

Rogers, M. L., Stanley, I. H., Hom, M. A., Chiurliza, B., Podlogar, M. C., & Joiner, T. E. (2016). Conceptual and empirical scrutiny of covarying depression out of suicidal ideation. Assessment. https://doi.org/10.1177/1073191116645907

Rohatgi, A. (2019). *WebPlotDigitizer* (4.2) [Computer software]. https://automeris.io/WebPlotDigitizer

Rufino, K. A., Marcus, D. K., Ellis, T. E., Boccaccini, M. T., Rufino, C., Marcus, K. A., Ellis, D. K., & Boccaccini, T. E. (2018). Further evidence that suicide risk is categorical: A taxometric analysis of data from an inpatient sample. Psychological Assessment. https://doi.org/10.1037/pas0000613

Schmaal, L., van Harmelen, A.-L., Chatzi, V., Lippard, E. T. C., Toenders, Y. J., Averill, L. A., Mazure, C. M., & Blumberg, H. P. (2019). Imaging suicidal thoughts and behaviors: A comprehensive review of 2 decades of neuroimaging studies. Molecular Psychiatry. https://doi.org/10.1038/s41380-019-0587-x

Silverman, M. M., Berman, A. L., Sanddal, N. D., O’Carroll, P. W., & Joiner, T. E. (2007). Rebuilding the tower of Babel: A revised nomenclature for the study of suicide and suicidal behaviors—Part 2: Suicide-related ideations, communications, and behaviors. Suicide and Life-Threatening Behavior, 37, 264–277. https://doi.org/10.1521/suli.2007.37.3.264

Smaldino, P. E., & McElreath, R. (2016). The natural selection of bad science. Royal Society Open Science, 3(9), 160384. https://doi.org/10.1098/rsos.160384

Song, W., Li, H., Guo, T., Jiang, S., & Wang, X. (2019). Effect of Affective Reward on Cognitive Event□related Potentials and its Relationship with Psychological Pain and Suicide Risk among Patients with Major Depressive Disorder. Suicide and Life-Threatening Behavior, 49(5), 1290–1306. https://doi.org/10.1111/sltb.12524

Tavakoli, P., Boafo, A., Dale, A., Robillard, R., Greenham, S. L., & Campbell, K. (2018). Event-Related Potential Measures of Attention Capture in Adolescent Inpatients With Acute Suicidal Behavior. Frontiers in Psychiatry, 9, 85. https://doi.org/10.3389/fpsyt.2018.00085

Tavakoli, P., Boafo, A., Jerome, E., & Campbell, K. (2020). Active and Passive Attentional Processing in Adolescent Suicide Attempters: An Event-Related Potential Study. Clinical EEG and Neuroscience, 155005942093308. https://doi.org/10.1177/1550059420933086

Tecce, J. J. (1972). Contingent negative variation (CNV) and psychological processes in man. Psychological Bulletin, 77(2), 73–108. https://doi.org/10.1037/h0032177

Thompson, P. M., Stein, J. L., Medland, S. E., Hibar, D. P., Vasquez, A. A., Renteria, M. E., Toro, R., Jahanshad, N., Schumann, G., Franke, B., Wright, M. J., Martin, N. G., Agartz, I., Alda, M., Alhusaini, S., Almasy, L., Almeida, J., Alpert, K., Andreasen, N. C., … Drevets, W. (2014). The ENIGMA Consortium: Large-scale collaborative analyses of neuroimaging and genetic data. Brain Imaging and Behavior, 8(2), 153–182. https://doi.org/10.1007/s11682-013-9269-5

Timsit, M., Koninckx, N., Dargent, J., & Dongier, O. F. M. (1970). Variations contingentes négatives en psychiatrie. Electroencephalography and Clinical Neurophysiology, 7.

Tsypes, A., & Gibb, B. E. (2020). Unpublished Data.

Tsypes, A., Owens, M., & Gibb, B. E. (2019). Blunted Neural Reward Responsiveness in Children With Recent Suicidal Ideation. Clinical Psychological Science, 216770261985634. https://doi.org/10.1177/2167702619856341

Tsypes, A., Owens, M., & Gibb, B. E. (2020). Reward Responsiveness in Suicide Attempters: An Electroencephalography/Event-Related Potential Study. Biological Psychiatry: Cognitive Neuroscience and Neuroimaging, S2451902220300963. https://doi.org/10.1016/j.bpsc.2020.04.003

van Boxtel, G. J. M., & Böcker, K. B. E. (2004). Cortical Measures of Anticipation. Journal of Psychophysiology, 18(2/3), 61–76. https://doi.org/10.1027/0269-8803.18.23.61

Van Heeringen, K., & Mann, J. J. (2014). The neurobiology of suicide. The Lancet Psychiatry, 1(1), 63–72. https://doi.org/10.1016/S2215-0366(14)70220-2

Van Orden, K. A., Witte, T. K., Cukrowicz, K. C., Braithwaite, S. R., Selby, E. A., & Joiner, T. E. (2010). The interpersonal theory of suicide. Psychological Review, 117(2), 575–600. https://doi.org/10.1037/a0018697

Vidal-Ribas, P., Janiri, D., Doucet, G. E., Pornpattananangkul, N., Nielson, D. M., Frangou, S., & Stringaris, A. (2021). Multimodal Neuroimaging of Suicidal Thoughts and Behaviors in a U.S. Population-Based Sample of School-Age Children. American Journal of Psychiatry, appi.ajp.2020.2. https://doi.org/10.1176/appi.ajp.2020.20020120

Viechtbauer, W. (2010). Conducting Meta-Analyses in R with the metafor Package. Journal of Statistical Software, 36(3). https://doi.org/10.18637/jss.v036.i03

Wasey, J. O. (2019). PRISMAstatement: Plot Flow Charts According to the “PRISMA” Statement. https://CRAN.R-project.org/package=PRISMAstatement

Weinberg, A., & Hajcak, G. (2010). Beyond good and evil: The time-course of neural activity elicited by specific picture content. Emotion, 10(6), 767–782. https://doi.org/10.1037/a0020242

Weinberg, A., & Hajcak, G. (2011). The late positive potential predicts subsequent interference with target processing. Journal of Cognitive Neuroscience, 23(10), 2994–3007. http://dx.doi.org.proxy.lib.fsu.edu/10.1162/jocn.2011.21630

Weinberg, A., May, A. M., Klonsky, E. D., Kotov, R., & Hajcak Greg. (2017). Decreased neural response to threat differentiates patients who have attempted suicide from nonattempters with current ideation. Clinical Psychological Science. https://doi.org/10.1177/2167702617718193

Weinberg, A., Perlman, G., Kotov, R., & Hajcak, G. (2016). Depression and reduced neural response to emotional images: Distinction from anxiety, and importance of symptom dimensions and age of onset. Journal of Abnormal Psychology, 125(1), 26–39. https://doi.org/10.1037/abn0000118

Wickham, H., Averick, M., Bryan, J., Chang, W., McGowan, L. D., François, R., Grolemund, G., Hayes, A., Henry, L., Hester, J., Kuhn, M., Pedersen, T. L., Miller, E., Bache, S. M., Müller, K., Ooms, J., Robinson, D., Seidel, D. P., Spinu, V., … Yutani, H. (2019). Welcome to the tidyverse. Journal of Open Source Software, 4(43), 1686. https://doi.org/10.21105/joss.01686

Wickham, H., & Bryan, J. (2019). readxl: Read Excel Files. https://CRAN.R-project.org/package=readxl

Witte, T. K., Holm-Denoma, J. M., Zuromski, K. L., Gauthier, J. M., & Ruscio, J. (2017). Individuals at high risk for suicide are categorically distinct from those at low risk. Psychological Assessment, 29(4), 382–393.

Woolf, S. H., & Schoomaker, H. (2019). Life Expectancy and Mortality Rates in the United States, 1959-2017. JAMA, 322(20), 1996. https://doi.org/10.1001/jama.2019.16932

World Health Organization. (2014). Preventing suicide: A global imperative. World Health Organization. https://doi.org/10.1002/9780470774120

